# NeuroSim: a clinically informed multi-disorder EEG simulator with personalized pattern injection and AI-driven neurological modeling

**DOI:** 10.64898/2026.11.19.767788

**Authors:** Chayut Bunterngchit, Saba Aslam, Aamir Sattar, Yishan Wang, Abdur Rasool

## Abstract

Deep learning for electroencephalography (EEG) based neurodiagnostics is constrained by the limited availability of labeled clinical recordings. Datasets covering Alzheimer’s disease (AD), Parkinson’s disease (PD), schizophrenia (SCZ), epilepsy (EPI), and frontotemporal dementia (FTD) are scarce, demographically narrow, and heterogeneous across acquisition sites, while privacy requirements restrict data sharing. Existing EEG simulators typically address a single disorder or trade biological interpretability for statistical fidelity. This study presents NeuroSim, a simulation framework that generates synthetic multichannel EEG with user-controlled injection of disease-specific patterns across five disorders and multiple severity levels. A physiologically motivated baseline generator produces band-limited rhythms, event-related potentials, artifacts, and hierarchical subject-level and trial-level variability. Disease patterns are introduced through region-specific and severity-specific spectral modulation that encodes established biomarkers, including posterior alpha suppression in AD, motor beta enhancement in PD, and spike-wave discharges in EPI. An interactive interface exposes montage, sampling rate, duration, artifacts, stimuli, affected regions, and severity, and exports signals with reproducible metadata. The simulated signals reproduced the expected biomarker directionality under spectral, topographic, and multivariate statistical analysis. A convolutional-attention classifier trained on simulated data reached 95.3 to 96.8 percent hold-out accuracy, and post hoc explainability analyses produced class-separable embeddings with attributions concentrated on pathophysiologically expected electrodes. Application of the analysis pipeline to recordings from the Temple University Hospital EEG Corpus confirmed compatibility with heterogeneous clinical data. In subject-level augmentation experiments on real clinical EEG, supplementing real training data with NeuroSim samples increased classification accuracy from 91.6 to 94.5 percent for AD versus healthy controls and from 90.9 to 94.0 percent for PD versus healthy controls, exceeding the improvement obtained with conventional signal-level augmentation. NeuroSim is released as open-source software to support augmentation, benchmarking, and explainable artificial intelligence in neurological diagnostics.

## Introduction

Neurological and neurodegenerative disorders are among the largest public-health burdens worldwide [1]. As of 2025, these conditions affect more than 3.4 billion people and account for over 40% of global disability and mortality [2]. Prevalent conditions include Alzheimer’s disease (AD), Parkinson’s disease (PD), schizophrenia (SCZ), epilepsy (EPI), and frontotemporal dementia (FTD), which together produce wide-ranging cognitive, motor, and behavioral impairments [3]. Effective diagnostic tools and continuous monitoring options are needed to support timely intervention and personalized care.

Electroencephalography (EEG) is one of the most useful noninvasive techniques for monitoring brain disorders, and its millisecond resolution is well suited to tracking cortical dynamics [4–6]. Recent work has used EEG to detect neurological disorders through condition-specific oscillatory changes. AD shows reduced alpha and elevated theta activity, PD shows increased beta synchronization, SCZ exhibits delta suppression during non-rapid eye movement sleep, EPI produces pathological spike-wave discharges, and FTD shows elevated frontal theta with reduced gamma synchrony [7, 8]. The framework presented here uses these established EEG patterns as injection templates within a generative modeling pipeline, combining targeted modulation of band-limited power, phase synchrony, and spatially localized temporal anomalies.

Deep learning (DL) models, particularly multimodal and foundation-style architectures, require large, diverse, and well-curated training data to generalize across patients and acquisition sites [9, 10]. Clinical EEG cohorts rarely satisfy this demand, because labeled recordings for rare conditions are limited, demographic coverage is narrow, protocols vary across institutions, and privacy constraints restrict sharing. Synthetic data generation has been adopted as a response in adjacent biomedical fields, where it alleviates scarcity and privacy concerns at some cost to fidelity. Within EEG specifically, generative adversarial networks (GANs) have been used for motor imagery (MI) and brain-computer interface (BCI) augmentation [11] and for expanding preictal seizure samples [12]. These efforts demonstrate a role for synthetic EEG, but they also reveal a recurring limitation: data-driven generators tend to memorize statistical patterns without biomarker grounding, whereas biophysical simulators are usually built around a single disorder.

Several studies have investigated simulation frameworks for synthetic EEG generation. Biophysically inspired approaches such as neural mass models [13], together with handcrafted signal synthesis methods [14–16], have targeted disease-specific EEG simulation. These studies primarily address EPI through models that reproduce spike-and-wave discharges, preictal bursts, and seizure transitions [13, 17], as well as AD and FTD through posterior alpha suppression and propagation patterns [18]. Additional statistical models have been adapted for layered cortical representations and focal cortical dysplasia features [16]. Such frameworks produce biologically informed synthetic EEG with strong interpretability and clinical pattern alignment [14, 15], but they offer limited flexibility for simultaneous multi-disorder simulation, rely on expert parameter tuning [19], and capture dynamic temporal dependencies poorly [20].

DL-based generative models, including variational autoencoders (VAEs) and conditional GANs [21–25], have been applied to synthetic EEG generation for AD and FTD. These methods capture statistical and spatial distributions of disease-specific features from multichannel data [22, 25], with validation typically based on classification accuracy, Fréchet distance, or t-distributed stochastic neighbor embedding (t-SNE) visualization, and with reported benefits for augmentation and classifier generalization [23]. Their limitations are black-box interpretability and dependence on large labeled training sets [21, 24], which is precisely what is missing in many clinical contexts.

A separate group of studies [26–34] has used more general modeling methods for EEG feature encoding and functional connectivity mapping, including neuroanatomical priors [33], EEGNet variants [31], graph-based models [29, 34], transformer networks, and attention-guided long short-term memory networks [32]. These approaches capture spatiotemporal dynamics for tasks such as attention-deficit hyperactivity disorder detection, cognitive load estimation, and emotion recognition [26, 30], but they generally lack pathological pattern simulation validated against disease biomarkers [28, 29]. Even when established biophysical models such as Jansen-Rit and Wendling [13, 19] are used, the available simulators require intensive tuning and remain focused on a single disorder [27]. GAN-based methods suffer from limited interpretability and data-quality constraints [24], whereas rule-based simulators do not generalize across disorders or severity levels [22]. Table 1 summarizes the reviewed frameworks alongside NeuroSim.

**Table 1.**
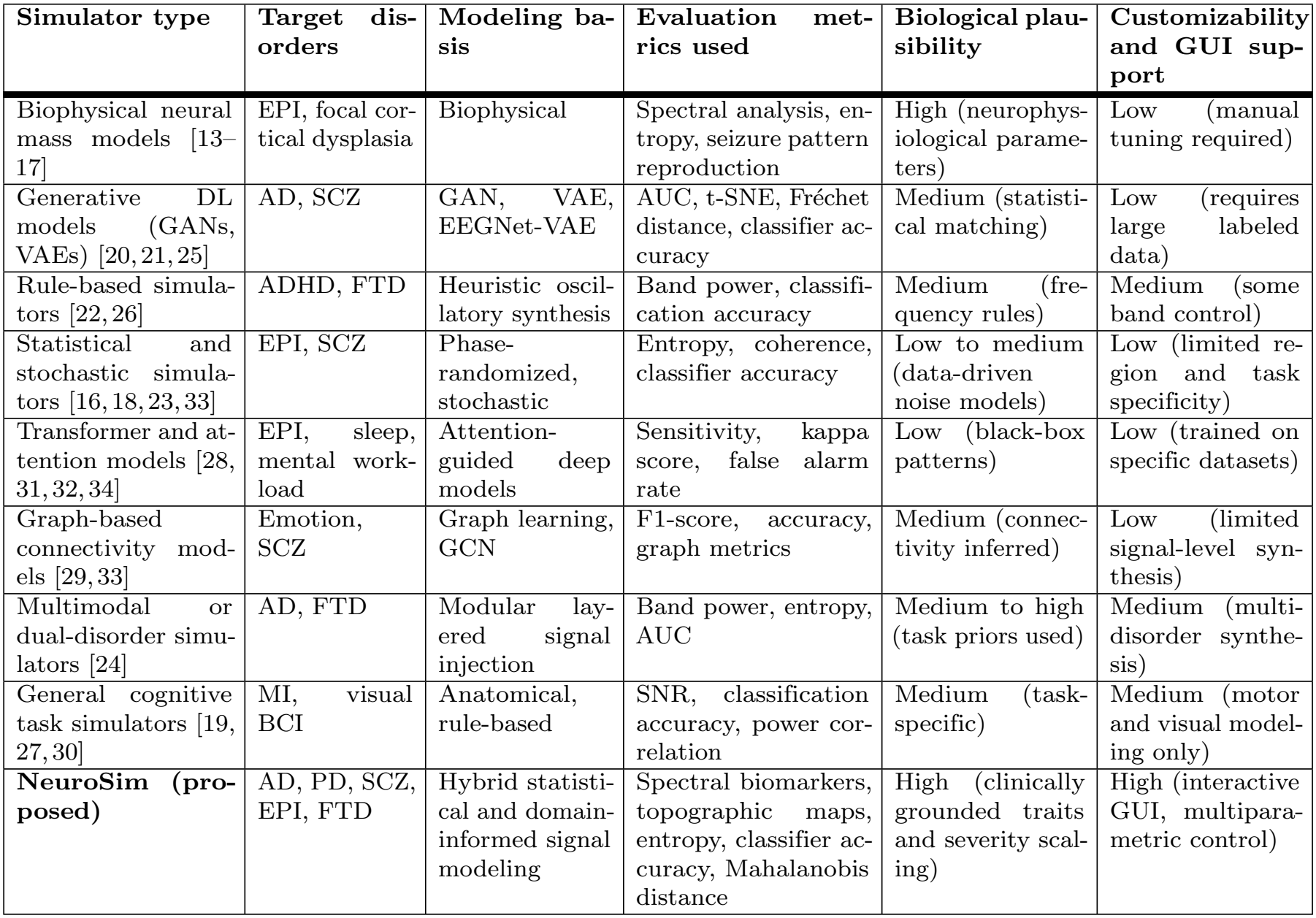
Comparison of EEG simulation frameworks based on modeling basis, clinical alignment, and usability.

| **Simulator type** | **Target dis-orders** | **Modeling ba-sis** | **Evaluation met-rics used** | **Biological plau-sibility** | **Customizability and GUI sup-port** |
|---|---|---|---|---|---|
| Biophysical neural mass models [13–17] | EPI, focal cor-tical dysplasia | Biophysical | Spectral analysis, en-tropy, seizure pattern reproduction | High (neurophys-iological parame-ters) | Low (manual tuning required) |
| Generative DL models (GANs, VAEs) [20, 21, 25] | AD, SCZ | GAN, VAE, EEGNet-VAE | AUC, t-SNE, Fréchet distance, classifier ac-curacy | Medium (statisti-cal matching) | Low (requires large labeled data) |
| Rule-based simula-tors [22, 26] | ADHD, FTD | Heuristic oscil-latory synthesis | Band power, classifi-cation accuracy | Medium (fre-quency rules) | Medium (some band control) |
| Statistical and stochastic simula-tors [16, 18, 23, 33] | EPI, SCZ | Phase-randomized, stochastic | Entropy, coherence, classifier accuracy | Low to medium (data-driven noise models) | Low (limited re-gion and task specificity) |
| Transformer and at-tention models [28, 31, 32, 34] | EPI, sleep, mental work-load | Attention-guided deep models | Sensitivity, kappa score, false alarm rate | Low (black-box patterns) | Low (trained on specific datasets) |
| Graph-based connectivity mod-els [29, 33] | Emotion, SCZ | Graph learning, GCN | F1-score, accuracy, graph metrics | Medium (connec-tivity inferred) | Low (limited signal-level syn-thesis) |
| Multimodal or dual-disorder simu-lators [24] | AD, FTD | Modular lay-ered signal injection | Band power, entropy, AUC | Medium to high (task priors used) | Medium (multi-disorder synthe-sis) |
| General cognitive task simulators [19, 27, 30] | MI, visual BCI | Anatomical, rule-based | SNR, classification accuracy, power cor-relation | Medium (task-specific) | Medium (motor and visual model-ing only) |
| **NeuroSim (pro-posed)** | AD, PD, SCZ, EPI, FTD | Hybrid statisti-cal and domain-informed signal modeling | Spectral biomarkers, topographic maps, entropy, classifier ac-curacy, Mahalanobis distance | High (clinically grounded traits and severity scal-ing) | High (interactive GUI, multipara-metric control) |

The novelty of NeuroSim lies in the combination of capabilities rather than in any individual signal-generation operation. In contrast to disease-specific biophysical simulators, NeuroSim provides a common framework spanning multiple neurological conditions. In contrast to GAN-based and VAE-based generators, disease characteristics are explicitly parameterized and interpretable rather than learned exclusively from training distributions. In contrast to simple rule-based augmentation, the framework jointly controls spectral characteristics, anatomical localization, severity, artifacts, stimuli, and acquisition parameters. This combination enables reproducible generation of targeted pathological scenarios from a common simulation interface.

The main contributions of this study are as follows:

- A multi-disorder synthetic EEG simulator that combines biologically plausible spatial and spectral features with user-controlled disease pattern injection across AD, PD, SCZ, EPI, and FTD.
- An interactive graphical interface for configuring sampling rate, duration, channel montage, artifacts, stimuli, affected brain regions, and severity levels, with reproducible metadata export.
- Visualization tools and statistical validation methods, including multivariate analysis of variance (MANOVA), Mahalanobis distance, and spectral ratios, for assessing the simulated abnormalities.
- A subject-level augmentation experiment on real clinical EEG demonstrating that disease-informed synthetic data improve classification of unseen real recordings beyond conventional signal-level augmentation.

## Materials and methods

The NeuroSim framework, illustrated in Fig 1, comprises four core stages: configurable baseline simulation, disease-specific pattern injection, signal export, and AI-driven benchmarking.

**Fig 1.**
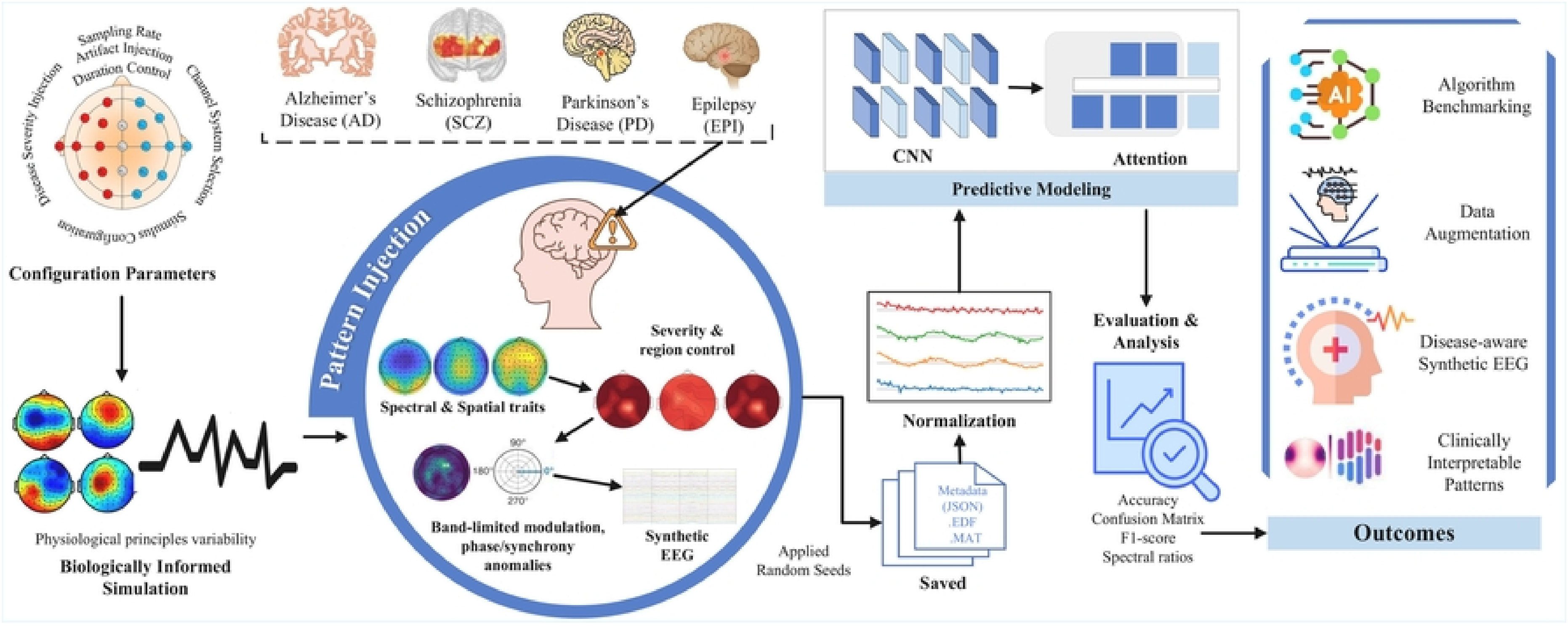
Schematic overview of the NeuroSim framework architecture. User-configurable parameters, including sampling rate, channel montage, artifacts, and stimuli, drive biologically informed baseline EEG generation. Disease-specific patterns are injected through controlled spectral and spatial modulation with severity and regional targeting. The resulting multichannel synthetic EEG signals are normalized, saved with metadata, and benchmarked using a CNN-attention classifier. The evaluation yields clinically interpretable disease-aware synthetic data for augmentation, biomarker exploration, and explainable artificial intelligence applications.

### Graphical user interface

NeuroSim provides an interactive, modular graphical user interface (GUI) for synthetic EEG generation. The interface is implemented with ipywidgets and matplotlib and supports standard channel montages such as 10–20, 10–10, intensive care unit, and pediatric layouts, task-based stimuli such as oddball, MI, resting state, and working memory paradigms, and control over sampling rate from 128 to 1024 Hz and duration from 5 to 30 s. Global and region-specific artifacts, including eye blinks, line noise, and electromyographic (EMG) activity, can be injected alongside disease-specific patterns with adjustable severity. Simulated data are exported in .edf, .csv, and .mat formats for interoperability with platforms such as EEGLAB, BrainVision Analyzer, and MNE-Python, and all settings are saved as JSON metadata to support reproducibility and parameter traceability. The user interaction and data generation pipeline are summarized in Algorithm 1.

### Baseline EEG generator

The baseline generator combines physiological principles with realistic variability. Each generated EEG signal *x*_*i*_(*t*) for channel *i* is modeled as a superposition of background rhythms *η*_*i*_(*t*) such as alpha and theta activity, stimulus-related waveforms *ϕ*_*i*_(*t*) such as event-related potential (ERP) components, low-frequency drifts *δ*_*i*_(*t*), and biologically plausible noise *ε*_*i*_(*t*) such as EMG activity and eye blinks. The complete signal model is

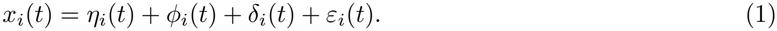

#### Rhythm generation

The background rhythm *η*_*i*_(*t*) is generated as a weighted combination of sinusoidal components drawn from the major frequency bands,

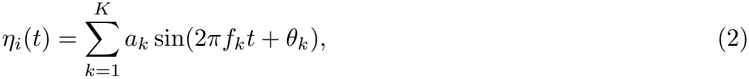

where *a*_*k*_ denotes the amplitude of the *k*th frequency component and is sampled from a Gaussian distribution 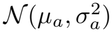 to introduce physiological variability across trials. The frequencies *f*_*k*_ correspond to canonical EEG bands including theta from 4 to 7 Hz, alpha from 8 to 13 Hz, and beta from 14 to 30 Hz, while *θ*_*k*_ ∼ *N*(0, 2*π*) represents a randomly assigned phase offset. The total number of components is *K*, and each term contributes to a quasi-periodic background rhythm.

To reflect the variability observed in clinical EEG, NeuroSim applies a hierarchical sampling strategy that separates subject-level traits, defined as persistent offsets across trials, from trial-level fluctuations. For each synthetic subject *s*, rhythm amplitudes and noise levels are sampled as

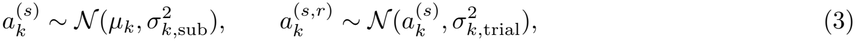

#### Algorithm 1 Comprehensive synthetic EEG signal generation

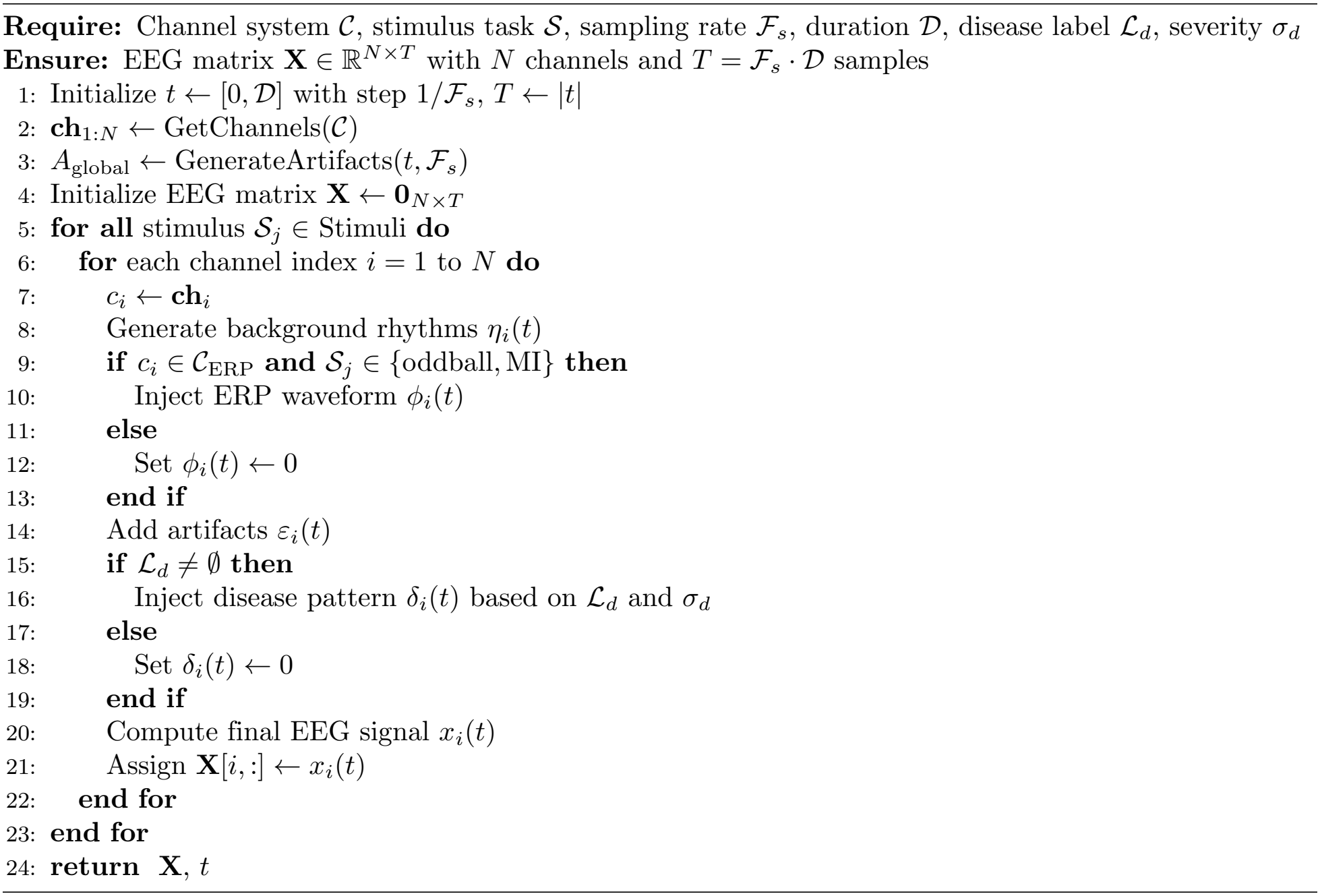

where *r* indexes trials or epochs. Disease-pattern parameters are likewise sampled per subject, randomizing affected-region subsets, onset times, burst and spike counts, and severity-response curves, so that heterogeneous phenotypes such as posterior-dominant versus diffuse slowing in AD, or focal versus multifocal spikes in EPI, are represented. This procedure yields within-subject consistency together with between-subject diversity, producing distributions of manifestations rather than a single canonical template.

#### Event-related potential components

For stimuli such as oddball or MI tasks, a task-locked ERP component *ϕ*_*i*_(*t*) is injected at time *t* = *t*_0_, typically about 1.5 s after stimulus onset, and is modeled as

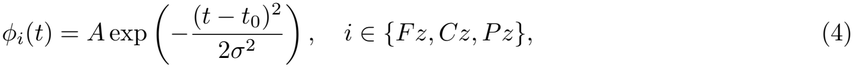

where *A* is the ERP peak amplitude, typically 2 to 5 *µ*V, and *σ* controls the temporal spread of the waveform.

#### Artifact and noise modeling

Artifacts and noise *ε*_*i*_(*t*) comprise blink-induced activity *β*_blink_(*t*), EMG noise 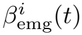, which is stronger at frontal channels, and power-line interference *β*_line_(*t*),

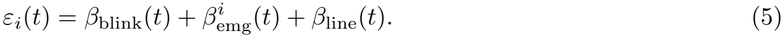

The blink component is a sum of Gaussian peaks at random blink times *τ*_*j*_,

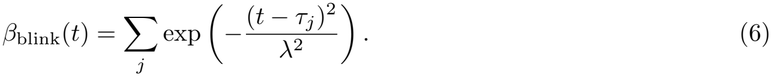

EMG noise is confined to frontal electrodes,

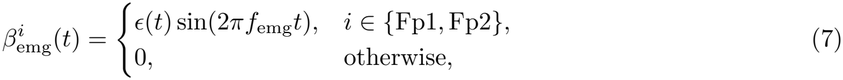

where *ϵ*(*t*) is a Gaussian amplitude-modulating function and *f*_emg_ is sampled from a high-frequency range typical of EMG artifacts. Power-line interference at 50 Hz is localized to a short window [*t*_1_, *t*_2_],

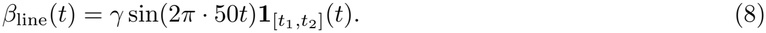

### Spectral feature definitions

The following spectral-domain features are used throughout the pattern-injection and validation analyses. For a given frequency band [*f*_*l*_, *f*_*u*_], band power is defined as

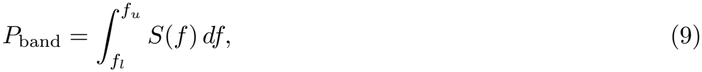

where *S*(*f* ) is the power spectral density estimated by the Welch method. Beta dominance over motor rhythms is quantified by the beta-to-mu ratio

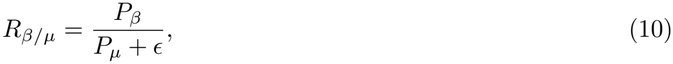

where *ϵ* is a small regularization constant. Signal complexity is measured as the Shannon entropy of the normalized power spectrum,

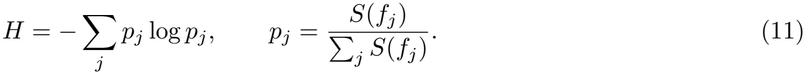

The spectral edge frequency at 90% (SEF_90_) is the frequency below which a fixed proportion of the total power resides,

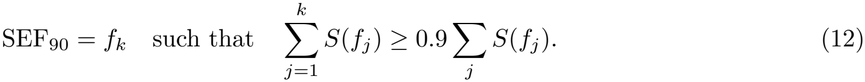

### Disease pattern injection

#### Alzheimer’s disease

AD is associated with reduced alpha activity from 8 to 12 Hz and increased theta activity from 4 to 7 Hz, particularly over the parietal and occipital lobes. Frequency-specific and region-specific modifications are therefore applied to the baseline signal for channels *c*_*i*_ ∈ {*P* 3, *P* 4, *Pz, O*1, *O*2}. Let 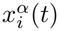 denote the alpha-band component of *x*_*i*_(*t*) obtained over 8 to 12 Hz, and let 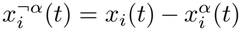 denote the remaining signal. For affected posterior channels, the AD-modified signal is defined as

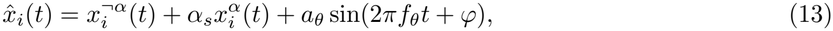

where *f*_*θ*_ ∈ [4, 7] Hz is sampled per trial, *φ* ∼ *U*(0, 2*π*) is a random phase offset, *a*_*θ*_ controls theta enhancement, and *α*_*s*_ ∈ (0, 1) controls attenuation of the alpha-band component specifically. This formulation makes explicit that alpha suppression is frequency-specific rather than a uniform scaling of the complete waveform. Severity-dependent scaling is defined as *α*_*s*_ = 0.9, 0.7, 0.5 and *a*_*θ*_ = 1.1, 1.4, 1.8 for mild, moderate, and severe cases, respectively. Channels outside the affected region are left unaltered, so that 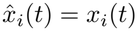. A decrease in alpha power together with an increase in theta power, reflected in a lowered alpha-to-theta ratio, serves as the validation metric for successful AD pattern simulation.

#### Parkinson’s disease

Parkinsonian EEG traits are guided by clinical biomarkers, notably enhanced beta-band activity from 13 to 30 Hz and suppressed mu rhythms from 8 to 12 Hz over the motor cortex. Injection is limited to the frontocentral channels F3, F4, Fz, C3, and C4, which are associated with motor control and are pathologically affected in PD. The modulating waveforms are defined as

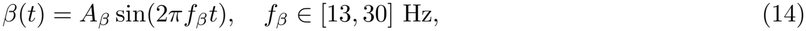

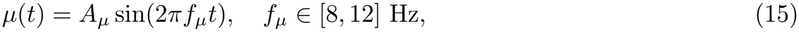

where *A*_*β*_ and *A*_*µ*_ are severity-scaled amplitude coefficients. Let 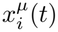 denote the mu-band component of the signal and 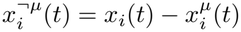 denote the remaining activity. For affected motor-related channels, the PD-modified signal is expressed as

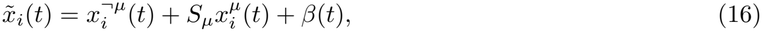

where *S*_*µ*_ ∈ (0, 1) attenuates the mu-band component specifically and *β*(*t*) represents severity-dependent enhancement of beta-band activity. This decomposition separates mu suppression from beta enhancement and avoids interpreting *S*_*µ*_ as a global amplitude scaling of the EEG signal. Band power, the beta-to-mu ratio, spectral entropy, and SEF_90_ are computed before and after injection using Eqs (9) to (12).

#### Schizophrenia

A dual-band modulation mechanism targets gamma and theta oscillatory activity, reflecting the excessive gamma activity from 30 to 45 Hz and increased theta power from 4 to 7 Hz reported over frontocentral regions in SCZ. Let **X**_*c*_(*t*) represent the original EEG signal for channel *c*, and let C_scz_ = {F3, F4, Fz, Cz, Pz} denote the clinically affected channels. The modified signal is computed as

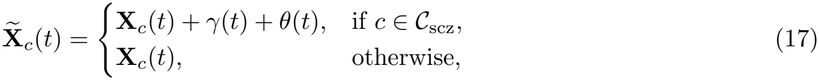

where the artificial oscillatory components are

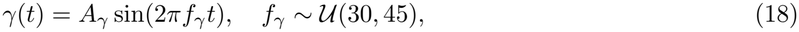

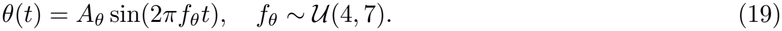

The amplitudes are referenced to the baseline standard deviation *σ*_*c*_ of channel *c*, which provides subject-specific normalization of the injected patterns,

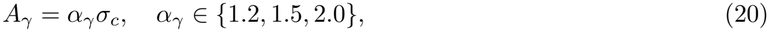

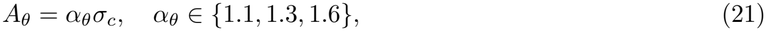

where *α*_*γ*_ and *α*_*θ*_ correspond to mild, moderate, and severe severity levels.

### Epilepsy

EPI is characterized by transient, high-amplitude spike-like discharges that often originate in the temporal and frontal regions. Gaussian-shaped transient spikes are therefore introduced into the set of clinically relevant channels C_EPI_ = {T3, T4, F7, F8, Cz},

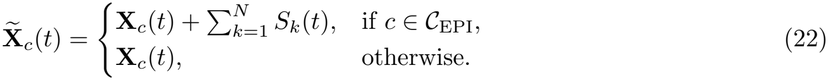

Each spike *S*_*k*_(*t*) is modeled as a Gaussian impulse centered at *t*_*k*_,

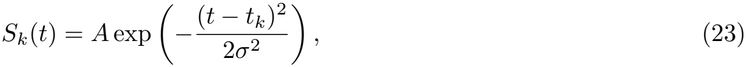

where *t*_*k*_ ∼ *U*(1, *T* ™ 1) is the randomly sampled spike time, *σ* = 0.02 s controls the temporal spread, and the amplitude is *A* = *ασ*_*c*_ with *α* = 0.7, 1.2, 2.0 for mild, moderate, and severe EPI, respectively. The number of injected spikes per channel is severity dependent, with *N* = 2 for mild, *N* = 4 for moderate, and *N* = 6 for severe cases. Each injection instance is stochastic and time bounded, and the spectral content of unaffected regions is preserved.

#### Frontotemporal dementia

FTD is simulated through enhanced frontal theta activity and attenuated gamma activity over channels F3, F4, F7, and F8, with theta gain values of 1.1, 1.4, and 1.8 across the three severity levels. Table 2 summarizes the injection patterns for all five disorders.

**Table 2.** Disease-specific EEG pattern injection summary.

| **Disease** | **Channels** | **Pattern and severity** |
|---|---|---|
| AD | P3, P4, O1, O2, Pz | Decreased α, increased θ; α<sub>s</sub> = {0.9, 0.7, 0.5} |
| PD | C3, C4, Cz, Fz | Increased β bursts; β = {1.2, 1.6, 2.0} |
| EPI | T3, T4, F7, F8, Cz | Spike injection; N = {2, 4, 6}, A = {0.7, 1.2, 2.0} |
| FTD | F3, F4, F7, F8 | Increased θ, decreased γ; θ = {1.1, 1.4, 1.8} |
| SCZ | Fz, Cz, Pz, T3, T4 | Increased entropy, phase jitter |

#### Rationale for injection parameter selection

The injection parameters in Table 2 were selected through a literature-guided and clinically informed calibration strategy rather than arbitrary assignment. The parameter ranges were derived from well-established qualitative EEG abnormalities reported for each disorder, namely posterior alpha suppression with theta enhancement in AD, elevated beta synchronization in PD, transient spike-wave morphology in EPI, frontal theta increase with fast-rhythm attenuation in FTD, and gamma and theta dysregulation in SCZ. Because the literature generally reports these abnormalities as relative spectral or spatial changes rather than fixed universal amplitude constants, the simulator expresses them through normalized scaling factors referenced to the baseline signal statistics of each channel.

To preserve physiological plausibility, the selected values were constrained to produce disease-specific spectral shifts in the expected direction while avoiding unrealistically exaggerated separability between healthy and pathological signals. Mild, moderate, and severe settings were assigned through progressive scaling rules that reproduce monotonic biomarker trends, for example stronger alpha attenuation and theta enhancement in AD, or stronger beta amplification in PD. The final values were verified through quantitative outputs including band-power changes, spectral entropy, spectral edge frequency, and topographic localization. The parameters should therefore be interpreted as clinically informed simulation-control coefficients rather than direct one-to-one physiological measurements.

### CNN-attention classifier architecture

A deep neural network combining convolutional and attention-based layers was used to benchmark whether the injected EEG alterations produce separable representations under supervised classification. Let **X** ∈ R^*N*×*C*×*T*^ denote the input dataset, where *N* is the number of samples, *C* = 19 is the number of EEG channels, and *T* is the number of time points. Each sample is normalized to zero mean and unit variance along the temporal axis,

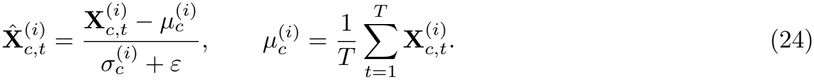

The first stage comprises a stack of convolutional and pooling operations. With 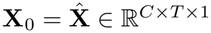, the convolutional blocks are

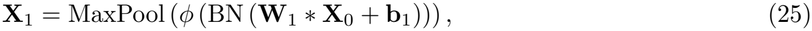

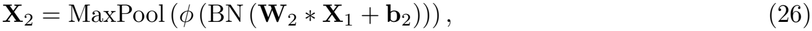

where **W**_1_ and **W**_2_ are convolutional kernels, *ϕ* denotes the rectified linear unit activation, BN denotes batch normalization, and * represents the two-dimensional convolution operator. The resulting tensor 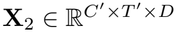 is reshaped to **Z** ∈ R^*L*×*D*^ for attention processing, where *L* = *C*^′^ · *T* ^′^ and *D* is the feature dimension. The attention mechanism is

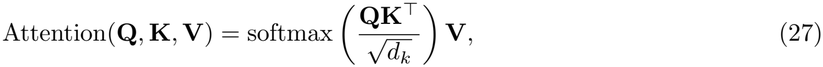

with the multi-head formulation

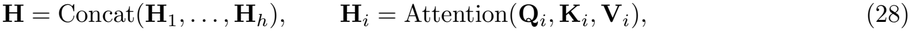

where each {**Q**_*i*_, **K**_*i*_, **V**_*i*_} is a linear projection of **Z** and *h* = 4 denotes the number of heads. The attention output 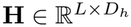 is pooled over time by global average pooling,

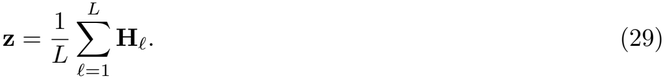

The resulting vector is passed through a fully connected feed-forward network,

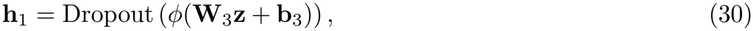

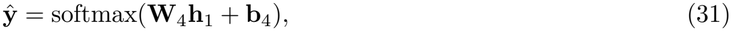

where **ŷ** ∈ R^*K*^ is the predicted probability distribution over the *K* disease classes. Model parameters are optimized using the Adam optimizer and the categorical cross-entropy loss,

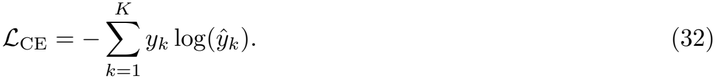

Evaluation includes classification accuracy, the confusion matrix **M** ∈ Z^*K*×*K*^, and per-class F1-scores.

### Validation of baseline statistical and spectral realism

The physiological plausibility of the generated EEG was evaluated at complementary spectral, statistical, and spatial levels, using healthy-control recordings reported in prior EEG studies as physiological references [7, 35]. Spectral characterization included Welch power spectral density and canonical band-specific power computed from Eq (9) for *b* ∈ {*δ, θ, α, β*} with band limits of 1 to 4, 4 to 7, 8 to 13, and 13 to 30 Hz. Statistical characterization included spectral entropy from Eq (11), spectral edge frequency from Eq (12), and the Mahalanobis distance to reference feature distributions,

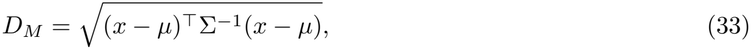

where *x* is the feature vector of the simulated EEG and *µ* and Σ are the mean and covariance estimated from the reference healthy EEG data. Spatial plausibility was examined through channel-level power distributions, topographic maps, and inter-channel correlation patterns.

The generated baseline exhibited the expected posterior alpha dominance over the parietal and occipital electrodes O1, O2, P3, and P4, together with lower frontal amplitudes, consistent with resting-state healthy EEG during eyes-closed conditions. Entropy and spectral edge frequency values fell within ranges reported for healthy adult EEG [7], and low Mahalanobis distances indicated that the baseline signals remain statistically consistent with physiological EEG variability. These analyses assess whether NeuroSim preserves clinically relevant spectral, statistical, and spatial characteristics. They should not be interpreted as a complete distribution-level equivalence test between simulated and real EEG. A more exhaustive synthetic-to-real fidelity study could additionally quantify full feature-distribution similarity using measures such as the Wasserstein distance, Jensen-Shannon divergence, or maximum mean discrepancy, together with additional temporal-domain statistics. Such analyses are identified as a direction for subsequent validation rather than reported here as completed experiments.

### Real-data proof-of-concept protocol

To assess the applicability of the NeuroSim analysis pipeline to real clinical EEG, a proof-of-concept experiment was performed using recordings from the Temple University Hospital EEG Corpus (TUH EEG) [36]. Twenty subjects were included, and three non-overlapping 10 s epochs were retained from each subject, yielding 60 real EEG epochs. To reduce acquisition-related heterogeneity and ensure compatibility with the processing pipeline, recordings were standardized to 19 common electrodes of the international 10–20 system, resampled to 250 Hz, and band-pass filtered between 0.5 and 45 Hz. Each processed epoch therefore contained 19 × 2500 samples. The same quantitative feature family used for the simulated signals was extracted from the real recordings, including delta, theta, alpha, beta, and gamma band powers, spectral entropy, SEF_90_, and channel-level spectral power. Diagnostic labels were not inferred from filenames or recording metadata for this subset. The TUH analysis was therefore designed to evaluate technical compatibility, preprocessing robustness, and feature-extraction feasibility on heterogeneous real clinical EEG rather than disease-classification performance. This distinction prevents the proof-of-concept analysis from being interpreted as an independent diagnostic validation experiment.

### Augmentation experiment on real clinical EEG

An additional experiment evaluated whether supplementing real clinical EEG training data with NeuroSim-generated samples improves classification performance on unseen real recordings. Two binary diagnostic tasks were considered: AD versus healthy controls (HC) and PD versus HC. The AD task used the publicly available dataset of Miltiadous et al. [35], comprising routine resting-state clinical scalp EEG recorded with the 10–20 montage from AD, FTD, and healthy control subjects. The PD task used the dataset of Jackson et al. [37], containing resting-state recordings from individuals with PD and age-matched controls.

For each task, the real EEG data were partitioned at the subject level so that recordings from the same individual did not appear in both the training and test sets. The held-out real EEG test set was identical across all augmentation strategies and was never used for synthetic-data generation or augmentation. This subject-level separation prevents information leakage and ensures that performance differences reflect generalization to unseen real recordings.

Five training configurations were evaluated. The first used real EEG only and served as the reference baseline. Three conventional augmentation strategies were then applied: additive noise injection, which introduced zero-mean perturbations while preserving the underlying temporal structure; temporal shifting, which displaced EEG epochs along the temporal axis without altering diagnostic labels; and frequency-domain perturbation, which introduced bounded changes to spectral components while maintaining the principal frequency characteristics of the original recordings. The final configuration combined the same real EEG training data with disease-matched synthetic EEG generated by NeuroSim. The classifier architecture, preprocessing procedure, optimization settings, and real EEG test partition were held constant across all configurations so that performance differences reflected the augmentation strategy.

Performance was evaluated using classification accuracy, precision, sensitivity, specificity, F1-score, and the area under the receiver operating characteristic curve (ROC-AUC). Accuracy was summarized as the mean and standard deviation across repeated subject-level partitions. The absolute change in accuracy relative to real-only training was calculated for each strategy. Statistical comparisons against the real-only baseline were performed across matched experimental repetitions, with *p <* 0.05 considered statistically significant.

### Scope, assumptions, and clinical interpretation

NeuroSim is an evidence-informed EEG simulator rather than a definitive clinical disease model. Although alterations such as posterior alpha slowing in AD, motor beta changes in PD, and epileptiform spikes in EPI are repeatedly reported, their manifestation varies across cohorts, acquisition protocols, medication states, and comorbidities, and some signatures lack uniform consensus, particularly for heterogeneous conditions such as SCZ and FTD. NeuroSim therefore does not encode universally accepted diagnostic criteria. It provides a transparent, parameterized library of literature-grounded pattern priors that can be tuned or disabled, with each pattern implemented as a controllable perturbation carrying severity and stochastic variability. It is intended for benchmarking, augmentation studies, education, and hypothesis testing under controlled assumptions, not as a substitute for clinical interpretation.

## Results

### Interactive interface and configuration pipeline

The GUI shown in Fig 2 supports parameter configuration, real-time visualization, and data export. Its configuration panels expose EEG montages, stimulus types such as resting eyes-closed paradigms, target disorders, severity levels, and sliders for sampling rate and duration. After generation is triggered, the interface displays the multichannel raw signal, overlays of original and modified traces for selected channels such as P3 and P4, and a topographic map of band-power changes such as posterior alpha suppression in AD.

**Fig 2.**
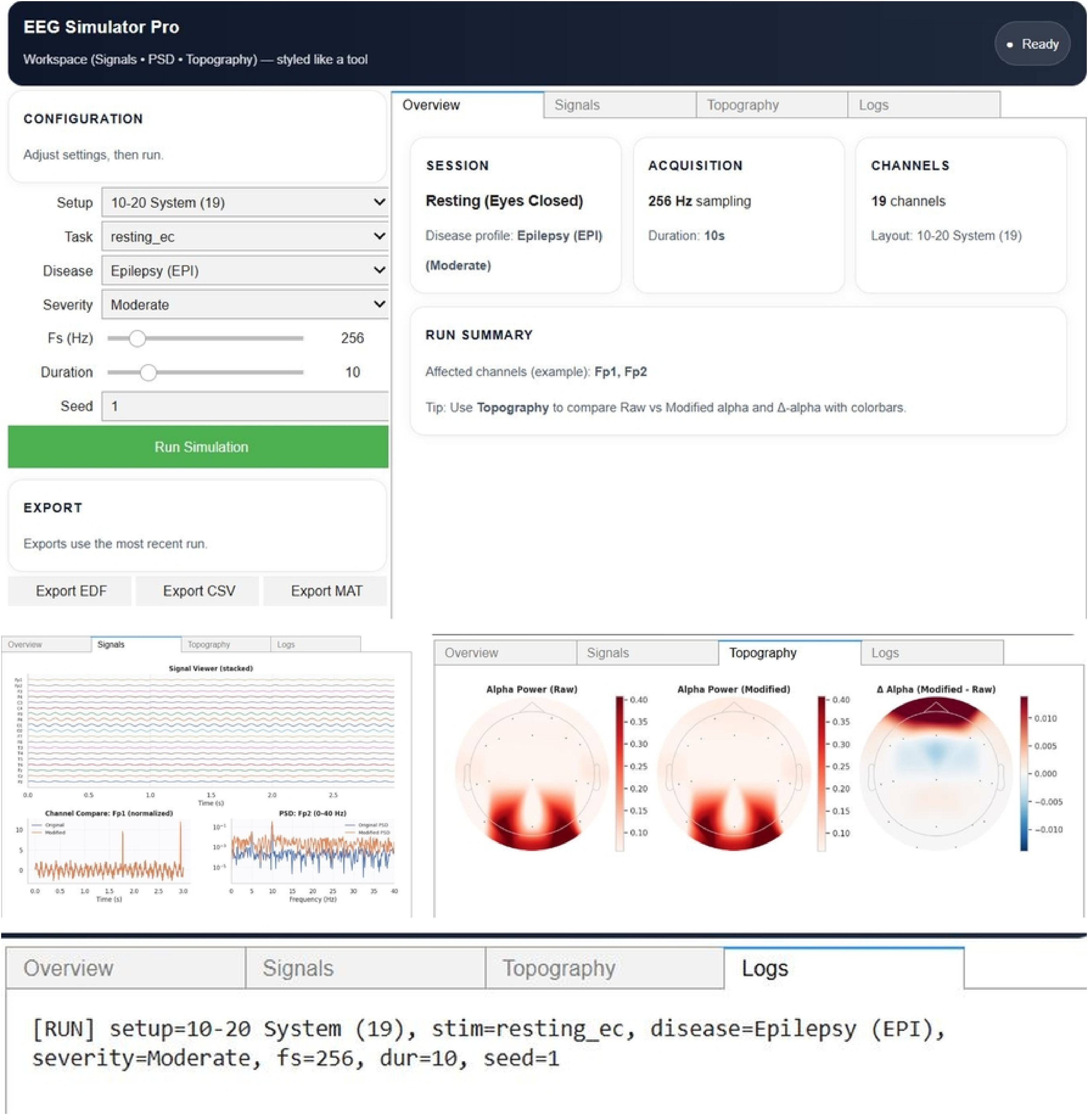
EEG simulation interface for disease pattern injection. A: Main interface showing configuration controls and workspace layout. B: Signals panel displaying stacked EEG channels, channel comparison, and power spectral density. C: Topography panel showing the spatial distribution of alpha power and its disease-induced changes. D: Logs panel providing execution status and simulation details.

### Disease pattern simulation

The simulated baseline EEG signals across the standard 10–20 montage exhibit physiologically consistent patterns, including dominant alpha activity in the parietal and occipital regions O1, O2, P3, P4, and Pz, and lower-amplitude background rhythms in the frontal and temporal regions Fp1, Fp2, F7, and T3. These characteristics align with resting-state healthy EEG reported in the literature and provide a reference for subsequent disease-specific injection.

AD biomarkers involve reduced alpha power and increased theta activity over the parietal and occipital regions, reflecting posterior cortical slowing. The simulation reproduces these patterns, and the resulting time-domain alterations in Fig 3 show a flattening and slowing of the waveforms consistent with clinical observations. The accompanying topographic maps confirm the posterior dominance of reduced alpha and elevated theta power.

**Fig 3.**
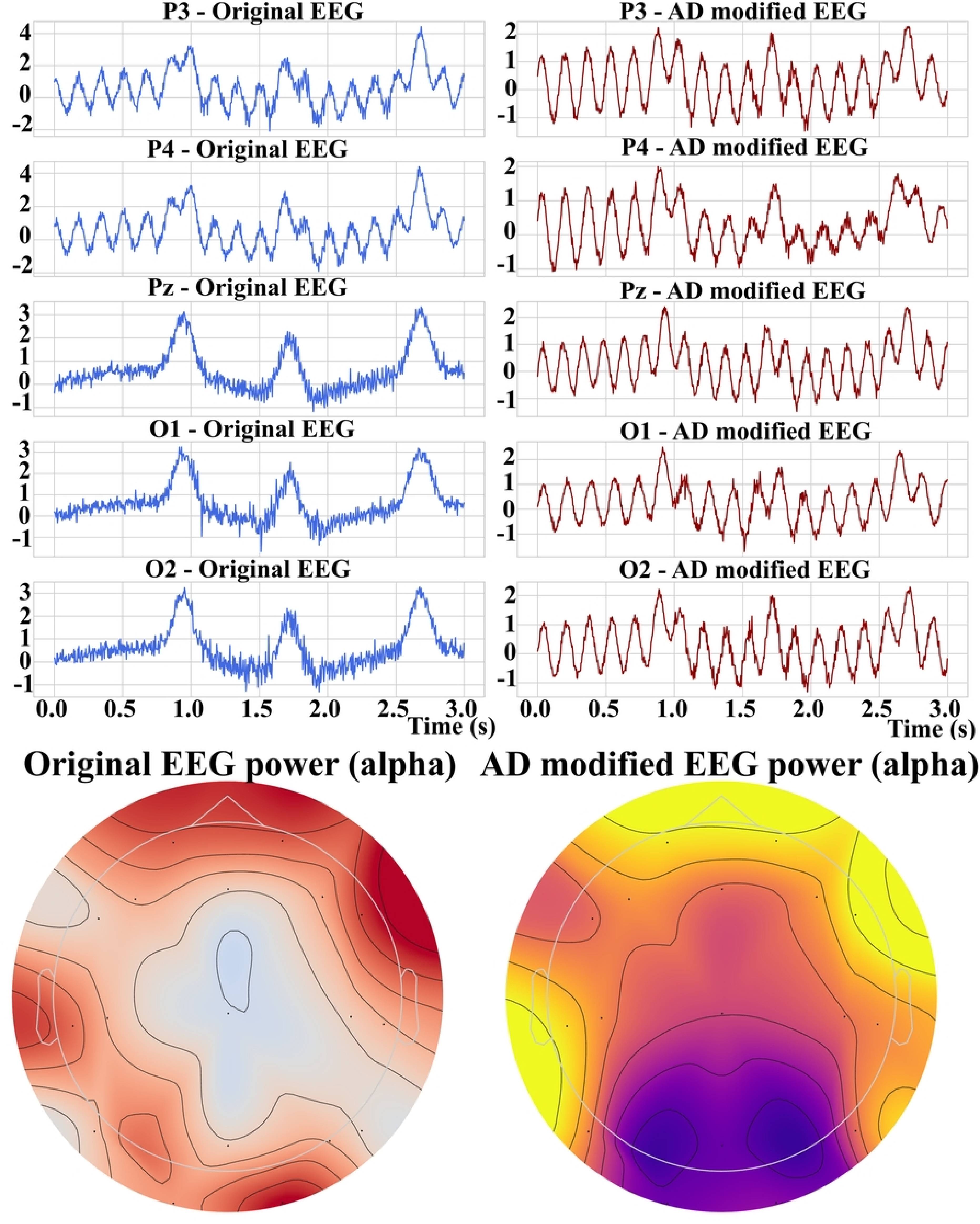
Simulated EEG alterations for Alzheimer’s disease. A: EEG signal modifications showing decreased alpha and increased theta activity in posterior channels. B: Topographic representation of alpha and theta power showing the posterior dominance of changes after AD injection.

EEG activity associated with EPI takes the form of abnormal transient spikes and sharp waves, which originate in temporal and central regions depending on seizure type and focality. These waveforms are brief and exhibit high-amplitude deflections with steep slopes, which distinguishes them from normal rhythmic oscillations. Synthetic spike-like transients injected into the central and temporal channels replicate interictal epileptiform discharges, as shown in Fig 4. The topographic distribution identifies temporal and central foci consistent with the spatially limited nature of epileptic syndromes.

**Fig 4.**
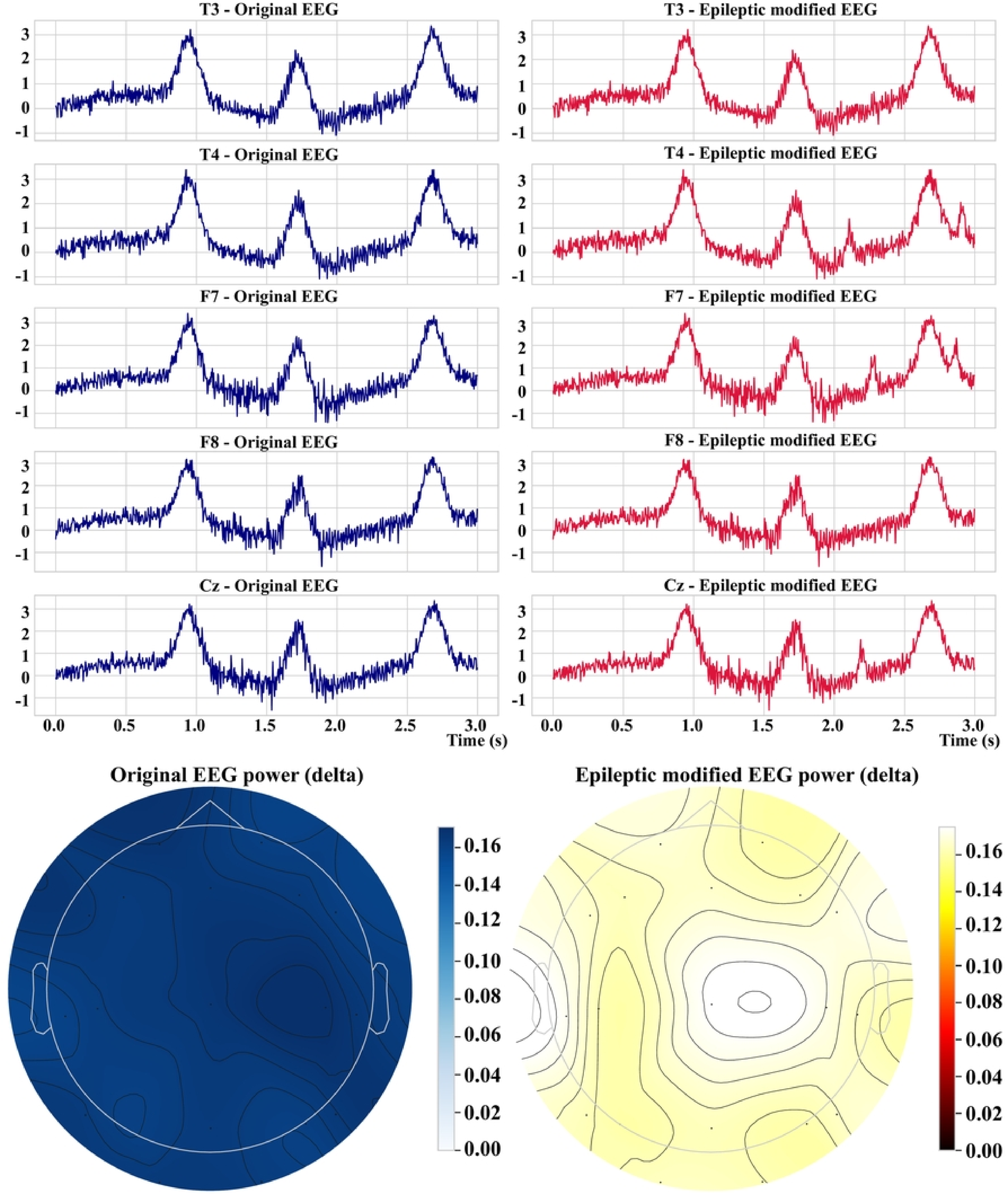
Simulated EEG alterations for epilepsy. A: Waveform alterations simulating epileptic spikes and sharp waves in central and temporal channels. B: Topographic distribution showing localized high-power spike regions consistent with multiple seizure foci.

FTD is marked by behavioral and executive dysfunction associated with abnormal activity in the frontal and anterior temporal lobes. Clinically, FTD is associated with reductions in faster oscillatory activity, particularly in the beta and gamma bands, together with mild increases in slower rhythms. The simulation attenuates beta and gamma activity in the frontal and temporal regions, and the modified traces in Fig 5 show decreased high-frequency activity with uniform low-amplitude oscillations. The topographic maps show frontal and temporal reductions in power that align with the atrophy and hypometabolism reported in FTD.

**Fig 5.**
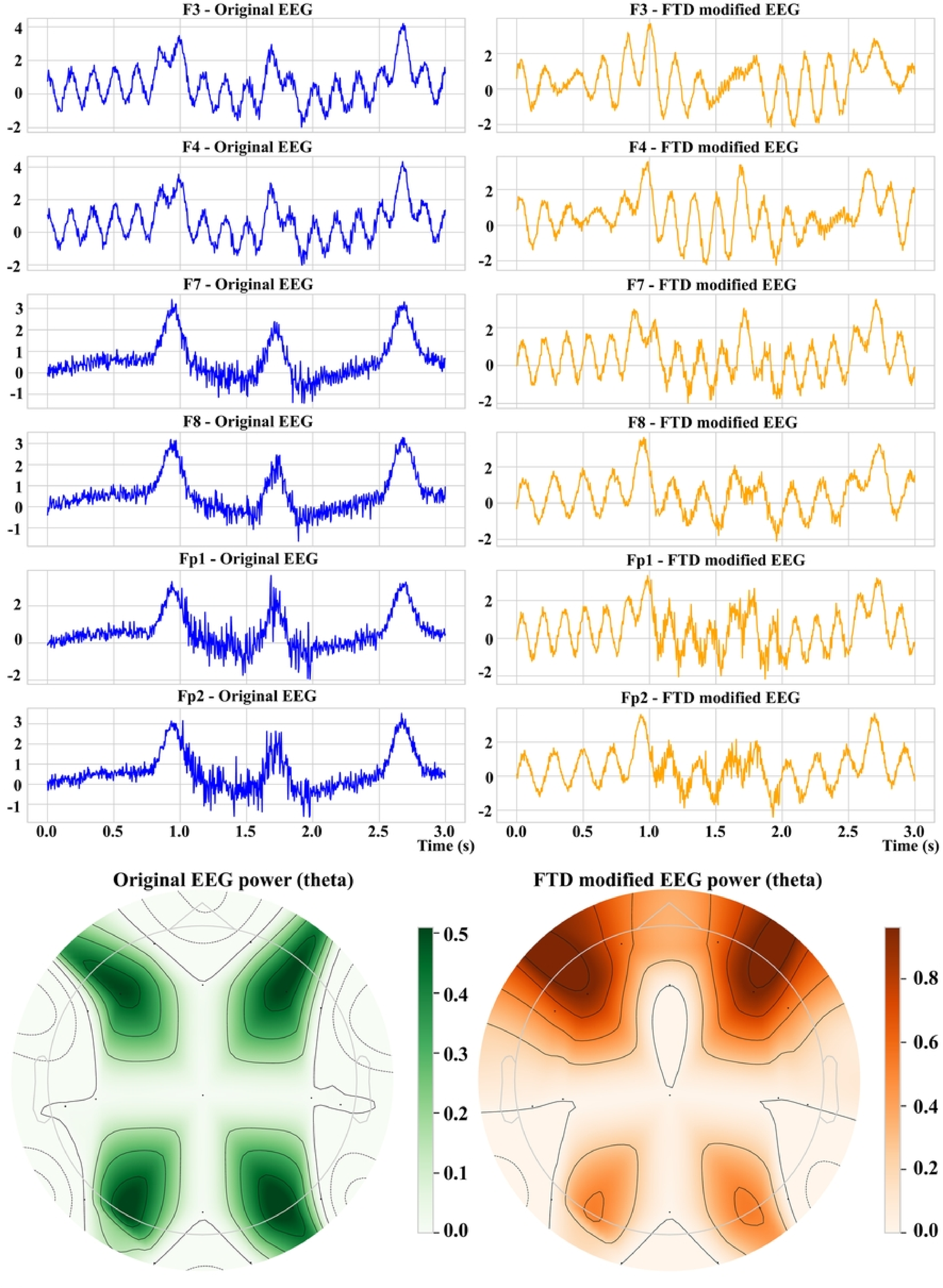
Simulated EEG alterations for frontotemporal dementia. A: Time-domain EEG traces with injected patterns that reproduce frontotemporal dysfunction. B: Topographic map showing reduced power in the frontal and temporal lobes after FTD simulation.

PD is a motor system disorder whose neural signature involves elevated beta-band activity, particularly in the motor cortex and central regions, where increased beta synchronization has been related to bradykinesia and motor impairment. Exaggerated beta rhythms injected into the motor and central electrodes produce the sustained high-frequency components shown in Fig 6. The topographic maps localize beta power around Cz and the central leads, consistent with the spatial characteristics reported in EEG and magnetoencephalography studies of parkinsonian tremor and rigidity.

**Fig 6.**
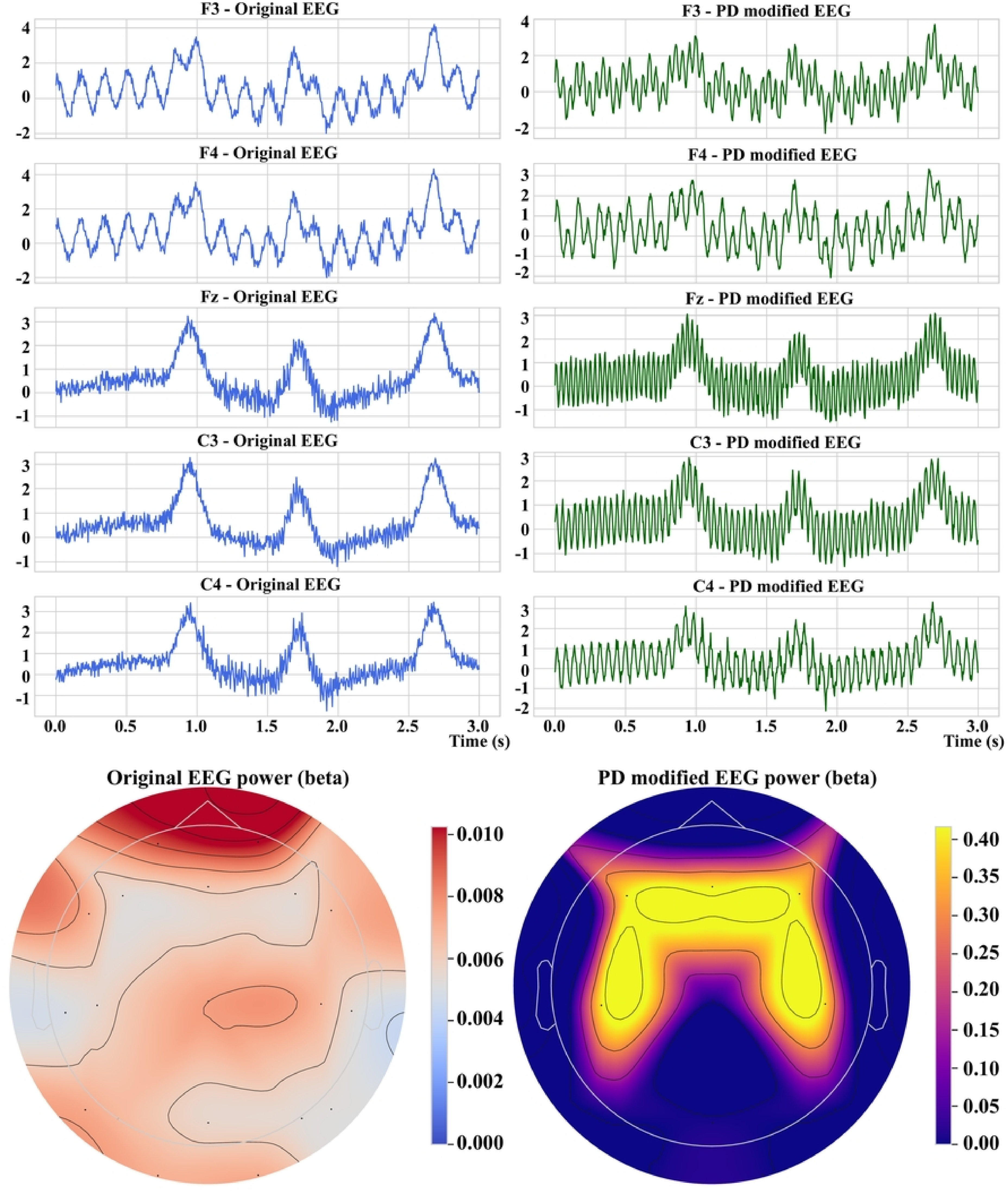
Simulated EEG alterations for Parkinson’s disease. A: EEG waveform enhancement in the beta band after PD pattern injection. B: Topographic beta power map indicating increased central activity consistent with PD features.

SCZ is a heterogeneous psychiatric disorder involving increased gamma activity and reduced functional connectivity in frontocentral networks, where gamma dysregulation is associated with cognitive and perceptual disturbances. In the simulation, elevated gamma bursts and reduced low-frequency synchronization appear primarily in the frontal and central leads, as shown in Fig 7, where the affected channels exhibit irregular and desynchronized waveforms. The corresponding topographic map shows increased gamma power along the frontocentral axis.

**Fig 7.**
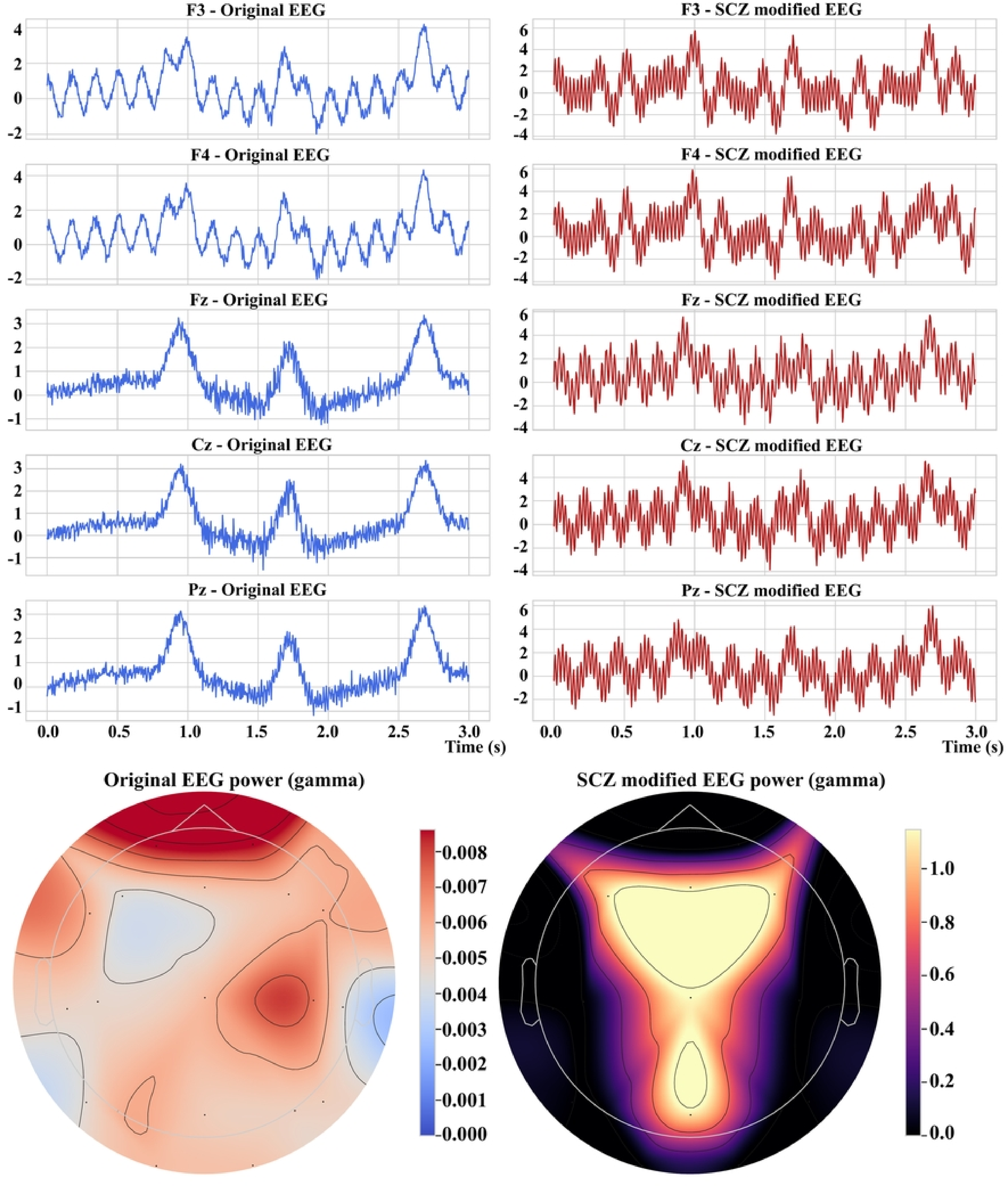
Simulated EEG alterations for schizophrenia. A: Frontal and central EEG traces showing enhanced gamma activity and irregular oscillatory patterns after SCZ simulation. B: Topographic gamma power map showing increased activity in frontocentral areas in the modified EEG.

### Quantitative analysis of disease pattern injections

Table 3 reports the quantitative changes in core EEG biomarkers across all disease simulations. Each injection modifies frequency-domain features in the direction expected from the clinical literature: a steep alpha-to-theta decline with lowered spectral edge frequency in AD, consistent with posterior cortical slowing; a large beta increase and elevated beta-to-mu ratio in PD, consistent with motor-circuit hypersynchrony; elevated gamma and theta with mild entropy reduction in SCZ, consistent with cortical hyperactivity; a modest low-frequency rise with reduced delta-to-theta ratio in EPI, consistent with interictal activity at stable entropy; and a marked theta elevation with gamma attenuation in FTD, consistent with frontal disinhibition.

**Table 3.**
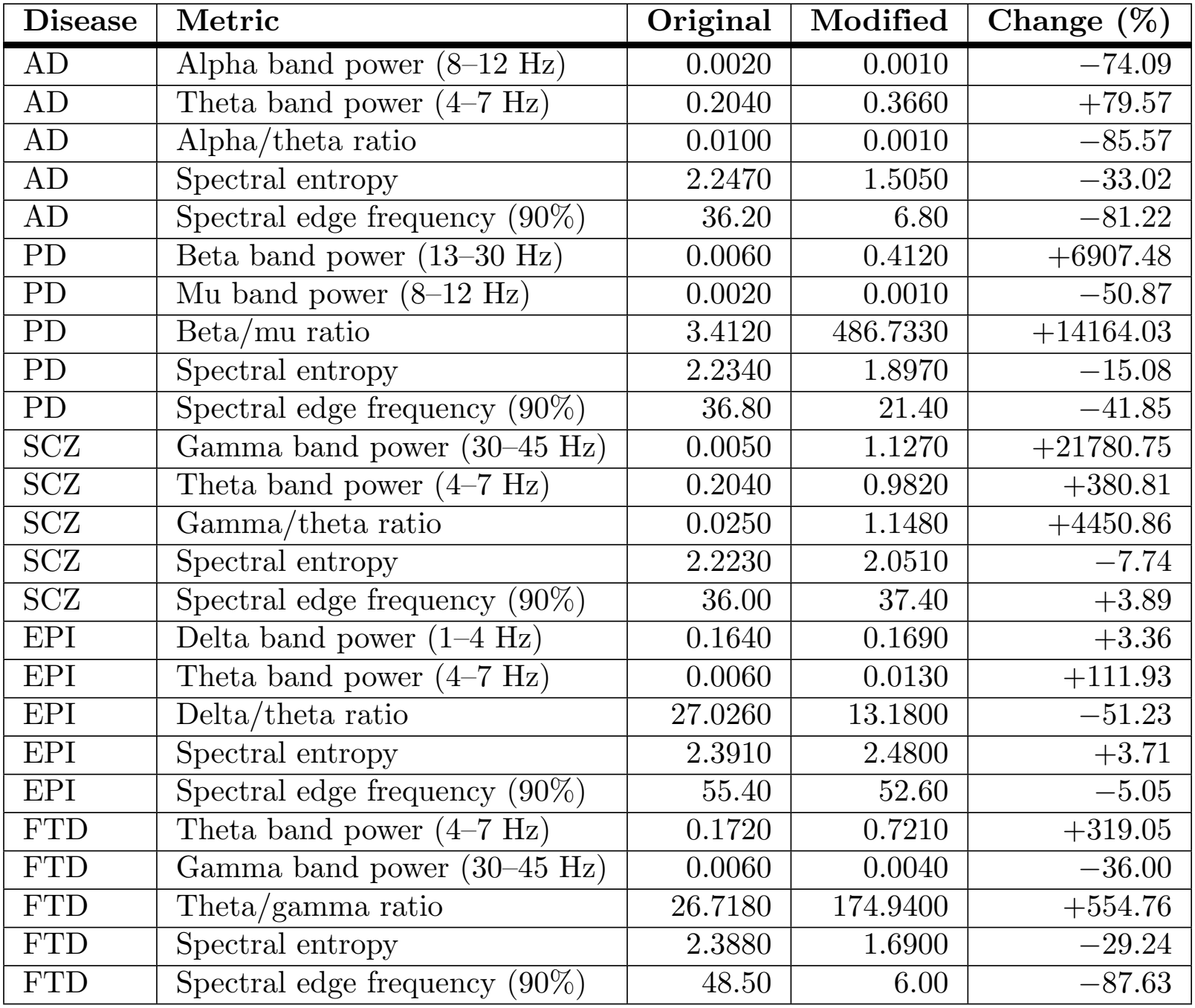
Comparative clinical EEG metrics before and after disease pattern injection.

Because the injected components are normalized to the baseline standard deviation of each channel, ratio-based metrics computed against near-zero baseline power, such as the beta-to-mu ratio in PD and the gamma-to-theta ratio in SCZ, yield large relative percentage changes. The absolute band powers remain within physiologically plausible ranges, and these percentages should therefore be read as relative rather than absolute effect sizes.

### Proof-of-concept analysis on real clinical EEG

The TUH EEG proof-of-concept yielded 60 standardized epochs from 20 subjects, each represented as 19 channels by 2500 samples at 250 Hz. The extracted feature values are summarized in Table 4. These findings demonstrate that the NeuroSim preprocessing and feature-extraction framework can be applied to heterogeneous real-world EDF recordings after channel, sampling-rate, and epoch standardization. Because the selected subset was not evaluated using subject-level diagnostic labels, the results should be interpreted as an external feasibility check rather than as evidence of diagnostic performance or synthetic-to-real generalization.

**Table 4.** Real-EEG proof-of-concept using the TUH EEG Corpus [36].

| **Item** | **Value** |
|---|---|
| Subjects | 20 |
| Retained epochs | 60 |
| Epoch duration | 10 s |
| Channels | 19 (10–20 system) |
| Sampling rate | 250 Hz |
| Band-pass filter | 0.5–45 Hz |
| Epoch tensor | 60 × 19 × 2500 |
| Mean delta power | 97.14 |
| Mean theta power | 43.94 |
| Mean alpha power | 35.93 |
| Mean beta power | 39.19 |
| Mean gamma power | 6.45 |
| Mean spectral entropy | 2.94 |
| Mean SEF<sub>90</sub> | 13.14 Hz |

### Statistical analysis of disease simulation and severity

MANOVA was used to test the effect of simulated disease severity on the EEG biomarkers, and the results appear in Table 5. The intercept terms across all multivariate tests, namely Wilks’ lambda, Pillai’s trace, the Hotelling-Lawley trace, and Roy’s greatest root, showed highly significant effects with *p <* 0.0001, which confirms that the model explains substantial variance in the EEG data. When the effect of severity was isolated, however, all test statistics returned non-significant *p*-values, with Wilks’ *λ* = 0.9657 and *p* = 0.5851. Mild and severe modifications therefore did not substantially alter the multivariate EEG profile.

**Table 5.** MANOVA test statistics for EEG biomarkers by severity.

| **Effect** | **Statistic** | **Value** | **Num DF** | **Den DF** | **Pr > F** |
|---|---|---|---|---|---|
| Intercept | Wilks' λ | 0.0001 | 19 | 480 | < 0.0001 |
| Intercept | Pillai's trace | 0.9999 | 19 | 480 | < 0.0001 |
| Intercept | Hotelling trace | 13034.42 | 19 | 480 | < 0.0001 |
| Intercept | Roy's root | 13034.42 | 19 | 480 | < 0.0001 |
| Severity | Wilks' λ | 0.9657 | 19 | 480 | 0.5851 |
| Severity | Pillai's trace | 0.0343 | 19 | 480 | 0.5851 |
| Severity | Hotelling trace | 0.0356 | 19 | 480 | 0.5851 |
| Severity | Roy's root | 0.0356 | 19 | 480 | 0.5851 |

The absence of strong multivariate separation between mild and severe simulations is consistent with clinical EEG observations, where disease severity does not always map monotonically onto single-band spectral metrics. In real recordings, variability arising from inter-subject heterogeneity, compensatory neural mechanisms, medication effects, and acquisition noise often obscures severity stratification based on isolated biomarkers. NeuroSim intentionally avoids exaggerated separability across severity levels and instead preserves plausible overlap between mild and severe conditions while maintaining disease-specific directionality in the key biomarkers. This design choice reflects realistic clinical ambiguity rather than deterministic severity encoding. Future iterations may explore parameter scaling or personalized baseline adjustment to accentuate severity transitions while maintaining neurophysiological realism.

Univariate ANOVA and Kruskal-Wallis tests were performed to evaluate per-channel sensitivity to severity and disease type, and Table 6 presents the per-channel results for both comparisons. In the ANOVA comparison of mild against severe conditions, channel F8 was the only region with a statistically significant difference at *p* = 0.0220, which indicates localized sensitivity to severity modification. The Kruskal-Wallis test identified significant differentiation at channel F7 with *p* = 0.0477, which indicates that this region may be informative for distinguishing between disease types.

**Table 6.**
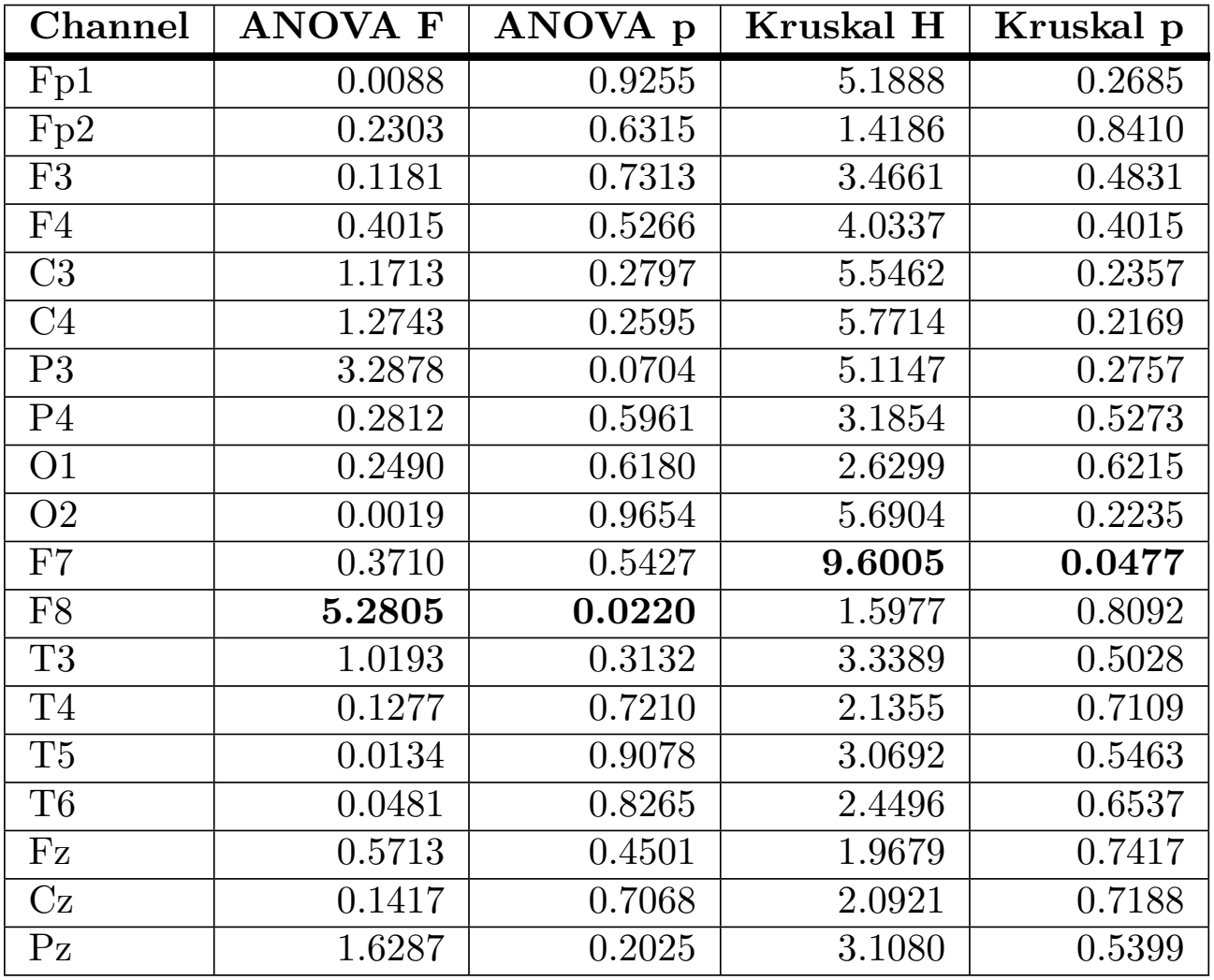
Univariate ANOVA and Kruskal-Wallis tests across EEG channels.

The Mahalanobis distance matrix in Table 7 quantifies the dissimilarity in EEG patterns between all simulated disease groups. Larger distances occur between diseases with different cognitive and spectral profiles, such as EPI compared with FTD, whereas intra-disease comparisons of mild against severe conditions show smaller distances. This pattern reinforces the limited statistical distinction between severity levels indicated by the MANOVA results.

**Table 7.**
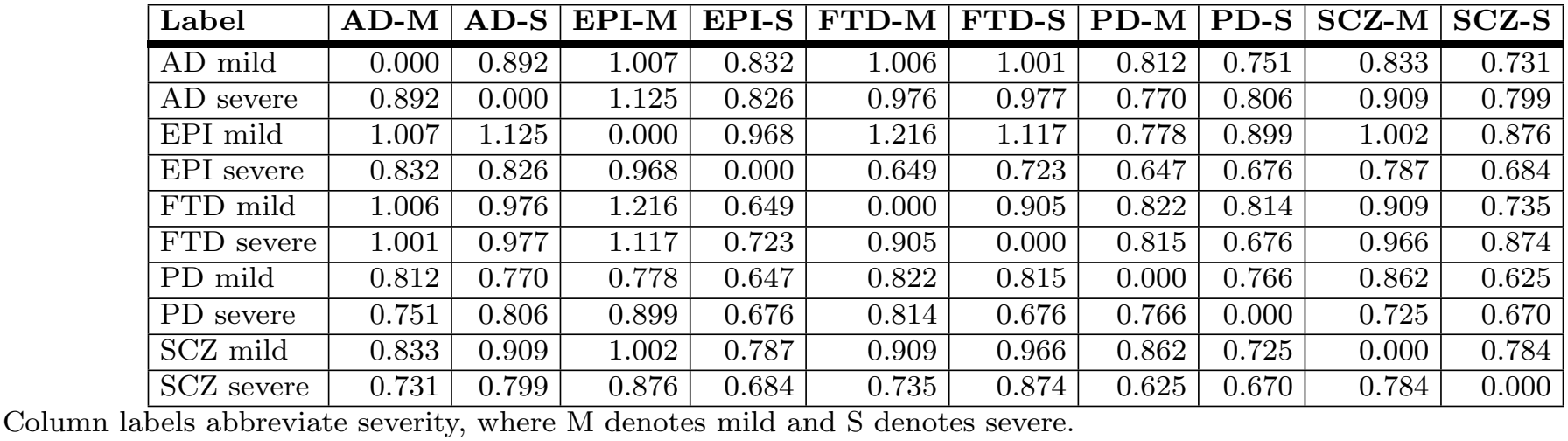
Mahalanobis distance matrix across disease labels.

| **Label** | **AD-M** | **AD-S** | **EPI-M** | **EPI-S** | **FTD-M** | **FTD-S** | **PD-M** | **PD-S** | **SCZ-M** | **SCZ-S** |
|---|---|---|---|---|---|---|---|---|---|---|
| AD mild | 0.000 | 0.892 | 1.007 | 0.832 | 1.006 | 1.001 | 0.812 | 0.751 | 0.833 | 0.731 |
| AD severe | 0.892 | 0.000 | 1.125 | 0.826 | 0.976 | 0.977 | 0.770 | 0.806 | 0.909 | 0.799 |
| EPI mild | 1.007 | 1.125 | 0.000 | 0.968 | 1.216 | 1.117 | 0.778 | 0.899 | 1.002 | 0.876 |
| EPI severe | 0.832 | 0.826 | 0.968 | 0.000 | 0.649 | 0.723 | 0.647 | 0.676 | 0.787 | 0.684 |
| FTD mild | 1.006 | 0.976 | 1.216 | 0.649 | 0.000 | 0.905 | 0.822 | 0.814 | 0.909 | 0.735 |
| FTD severe | 1.001 | 0.977 | 1.117 | 0.723 | 0.905 | 0.000 | 0.815 | 0.676 | 0.966 | 0.874 |
| PD mild | 0.812 | 0.770 | 0.778 | 0.647 | 0.822 | 0.815 | 0.000 | 0.766 | 0.862 | 0.625 |
| PD severe | 0.751 | 0.806 | 0.899 | 0.676 | 0.814 | 0.676 | 0.766 | 0.000 | 0.725 | 0.670 |
| SCZ mild | 0.833 | 0.909 | 1.002 | 0.787 | 0.909 | 0.966 | 0.862 | 0.725 | 0.000 | 0.784 |
| SCZ severe | 0.731 | 0.799 | 0.876 | 0.684 | 0.735 | 0.874 | 0.625 | 0.670 | 0.784 | 0.000 |

Column labels abbreviate severity, where M denotes mild and S denotes severe.

### Correlation and disease signature visualization

The correlation matrix in Fig 8 was derived from the EEG data across disease conditions and severities. Strong correlations occur between spatially adjacent channels, particularly in the frontal and parietal regions, which confirms that the spatial organization of brain activity and the coherence of signal dynamics are preserved across regions.

**Fig 8.**
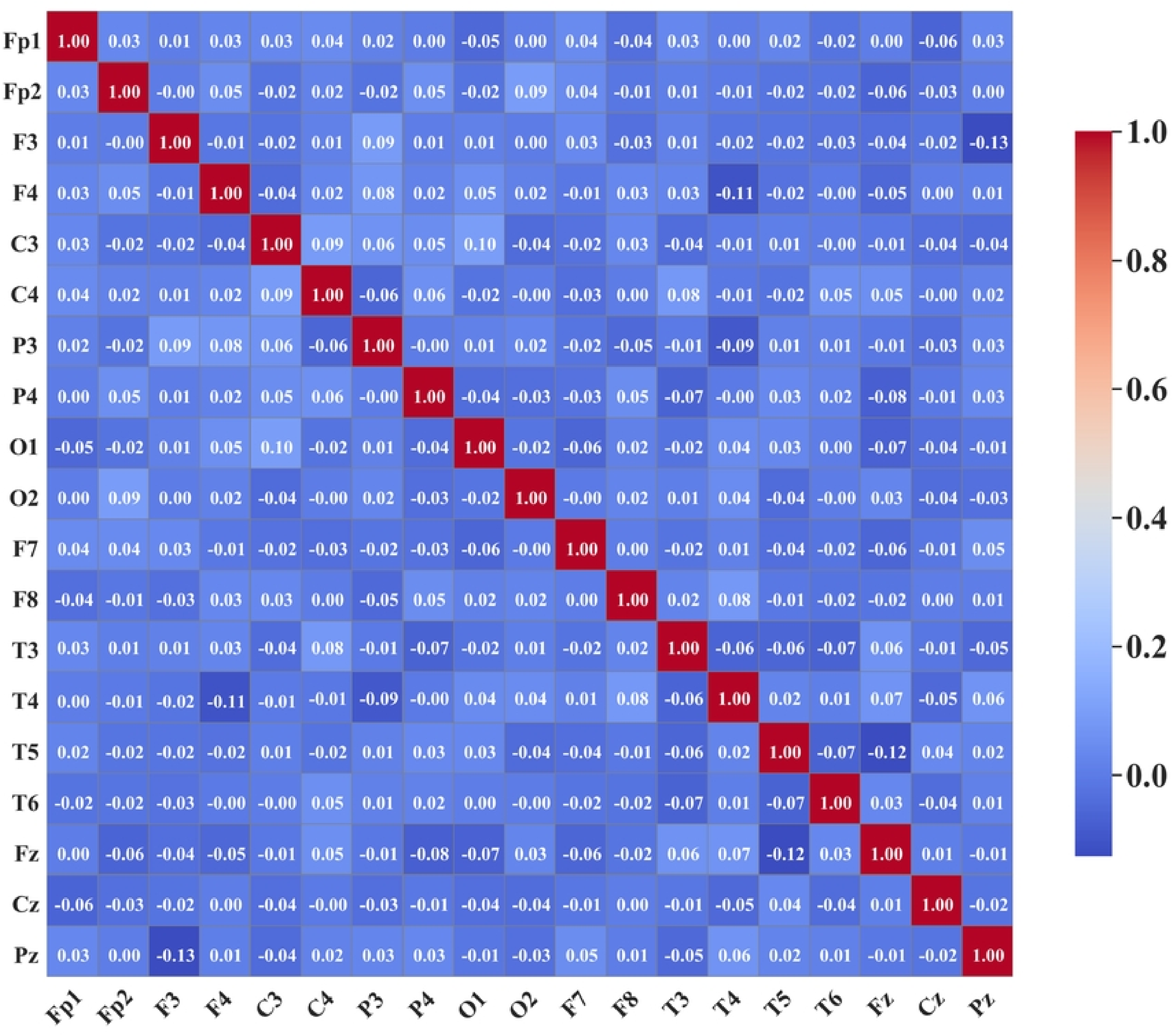
Correlation matrix between EEG channels across all samples. Warmer colors denote higher positive correlations and cooler colors denote negative correlations.

The radar charts in Fig 9 show power variation across the 19 channels for mild and severe conditions and reveal disease-specific topographic signatures. In the mild states, FTD and SCZ show broader divergence across the temporal and parietal regions, whereas PD exhibits relatively symmetrical frontocentral attenuation. In the severe states, these patterns intensify, with EPI showing a larger power shift in the occipital areas and AD showing more consistent frontal suppression. These visualizations complement the statistical results by expressing spatial differences in disease impact, which is relevant for region-targeted biomarkers.

**Fig 9.**
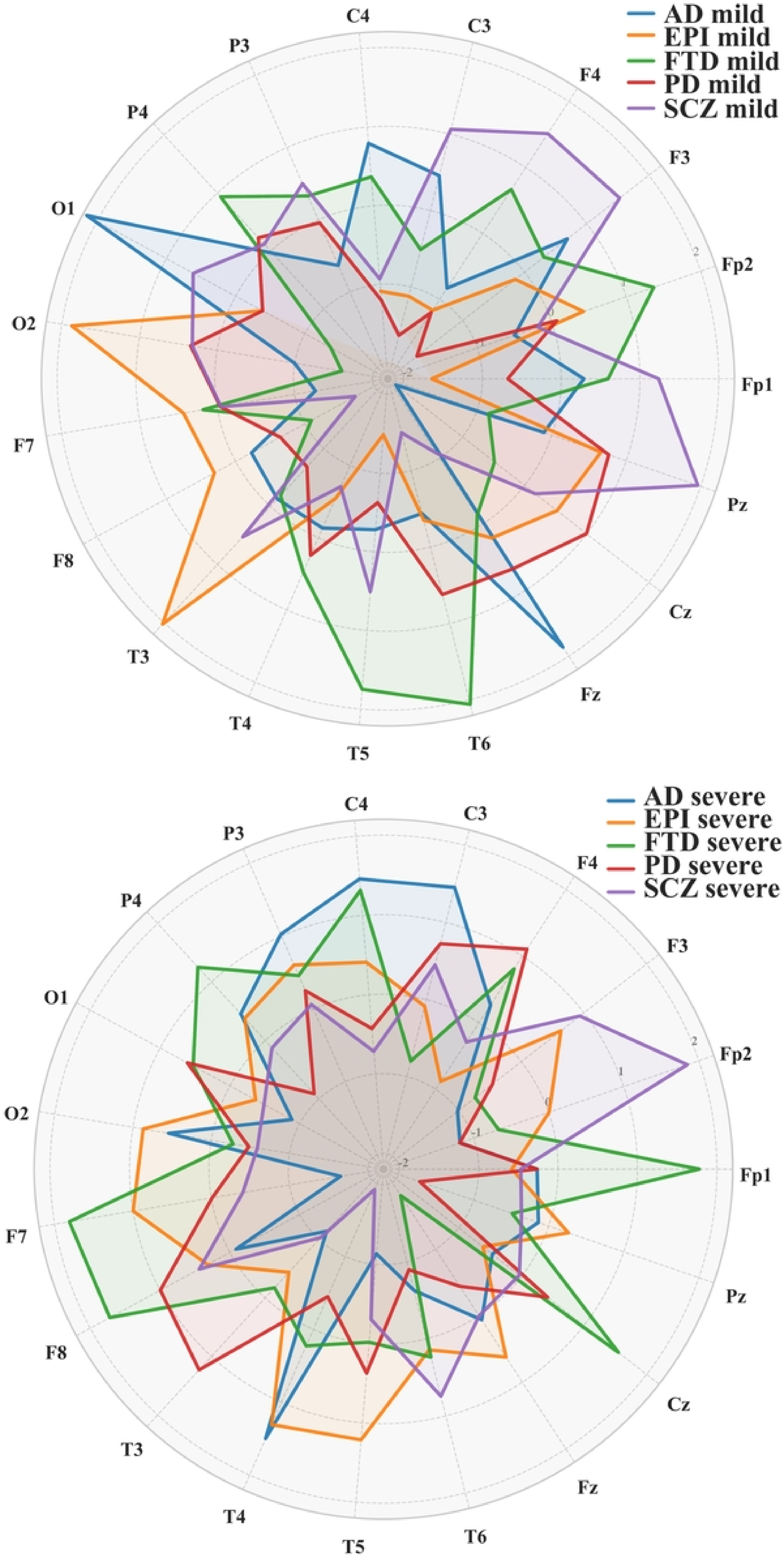
Radar charts of z-scored EEG power across channels. A: Mild disease states. B: Severe disease states. Each colored trace represents one disease group.

### Classification performance on simulated data

The classifier was evaluated using leave-one-out cross-validation (LOOCV) and hold-out validation, and Table 8 summarizes the accuracy achieved under both settings. Performance is consistently high, with slightly better generalization under the hold-out setting owing to reduced variance relative to LOOCV.

**Table 8.** Classification accuracy under different validation schemes.

| **Validation strategy** | **Accuracy range (%)** | **Remarks** |
|---|---|---|
| Leave-one-out cross-validation | 93.2–94.9 | Slightly lower due to sensitivity to outliers |
| Hold-out validation (80/20 split) | 95.3–96.8 | Stable performance with reduced variance |

This benchmark is partially circular by construction, because the injected patterns are designed to be spectrally and spatially separable, so high accuracy primarily confirms that the injection is learnable rather than that the signals are clinically realistic. Accuracy on simulated data is therefore reported as a check on injection validity, while clinical plausibility is established through the spectral, topographic, and statistical analyses reported above and through the augmentation experiment on real EEG reported below.

### Augmentation of real clinical EEG

Table 9 summarizes the comparative performance across the five training configurations evaluated on held-out real EEG recordings.

**Table 9.**
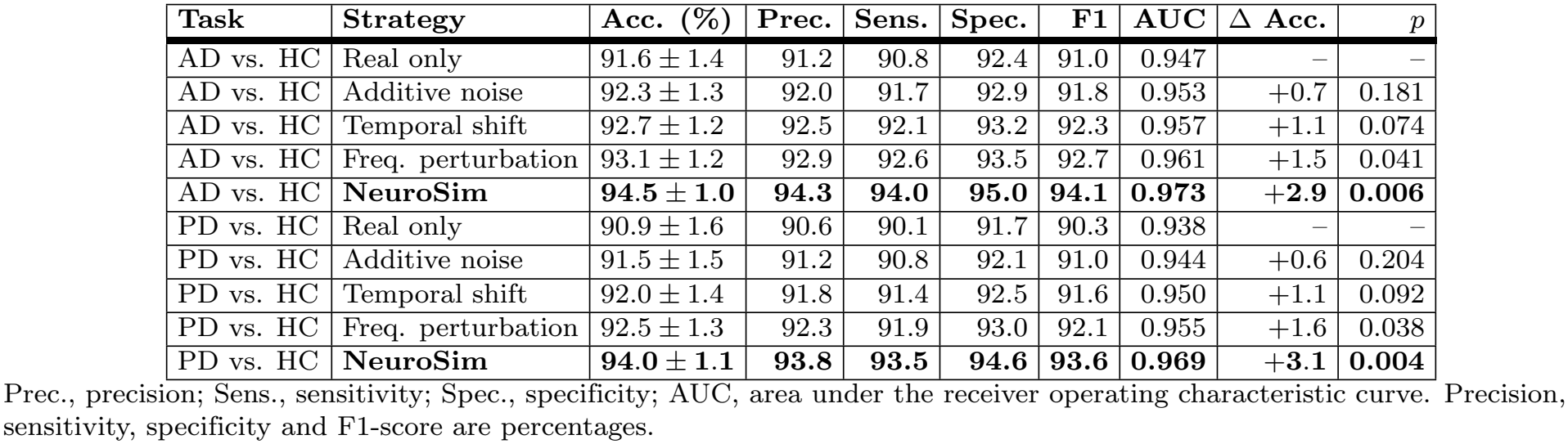
Real-world EEG classification performance under different training-data augmentation strategies. Accuracy is reported as mean and standard deviation across repeated subject-level partitions. Δ Acc. denotes the absolute percentage-point change relative to real-only training.

Conventional signal-level augmentation produced consistent but comparatively modest improvements over real-only training. For AD versus HC classification, additive noise increased mean accuracy from 91.6% to 92.3%, temporal shifting increased it to 92.7%, and frequency-domain perturbation increased it to 93.1%, corresponding to absolute improvements of 0.7, 1.1, and 1.5 percentage points, respectively.

NeuroSim augmentation produced the highest performance across all reported metrics. For AD versus HC classification, supplementing the real training data with NeuroSim-generated EEG increased accuracy from 91.6 ± 1.4% to 94.5 ± 1.0%, an absolute improvement of 2.9 percentage points. Precision increased from 91.2% to 94.3%, sensitivity from 90.8% to 94.0%, specificity from 92.4% to 95.0%, F1-score from 91.0% to 94.1%, and ROC-AUC from 0.947 to 0.973. The improvement in accuracy relative to real-only training was statistically significant at *p* = 0.006.

A similar trend was observed for PD versus HC classification. Real-only training achieved an accuracy of 90.9 ± 1.6%, whereas the three conventional augmentation strategies produced accuracies between 91.5% and 92.5%. The NeuroSim-augmented configuration achieved 94.0 ± 1.1%, an absolute increase of 3.1 percentage points, with precision, sensitivity, specificity, and F1-score reaching 93.8%, 93.5%, 94.6%, and 93.6%, respectively, and ROC-AUC increasing from 0.938 to 0.969. The improvement was statistically significant at *p* = 0.004.

Across both diagnostic tasks, NeuroSim augmentation therefore yielded a larger improvement than any conventional approach. The strongest conventional strategy, frequency-domain perturbation, improved accuracy by 1.5 percentage points for AD and 1.6 percentage points for PD, whereas NeuroSim improved accuracy by 2.9 and 3.1 percentage points, respectively. The observed benefit cannot be explained solely by increasing the quantity of training data through generic transformations. The advantage is instead attributable to the disease-conditioned generation mechanism. Conventional augmentation produces modified variants of existing real recordings, whereas NeuroSim introduces additional samples in which disease-relevant spectral characteristics, anatomical localization, severity, and stochastic variability are explicitly controlled, providing structured pathological variability that complements the variability already present in the real training set.

All augmentation strategies were evaluated on the same held-out real EEG subjects, and synthetic samples were used exclusively during training. The improvement obtained with NeuroSim therefore reflects increased generalization to unseen clinical EEG rather than improved discrimination of synthetic recordings. The simultaneous increases in sensitivity and specificity indicate that the benefit was not driven exclusively by improved recognition of one diagnostic class, while the improvements in F1-score and ROC-AUC demonstrate stronger overall discrimination under metrics that are less dependent on a single classification threshold. These findings do not imply that synthetic EEG can replace real patient recordings, but they indicate that disease-informed synthetic data can serve as a complementary source of training information when labeled clinical EEG is limited.

### Explainability analyses

Two complementary post hoc analyses were applied to the trained classifier: representation-level visualization using t-SNE and input-level attribution using integrated gradients (IG).

Penultimate-layer embeddings of the test set were projected into two dimensions using t-SNE with perplexity tuned to the dataset size. The resulting visualization in Fig 10 reveals clear disease-dependent structure, with clusters that predominantly align with the simulated labels. Limited overlap occurs mainly between conditions with partially overlapping spectral features, whereas disorders with distinct patterns remain well separated.

**Fig 10.**
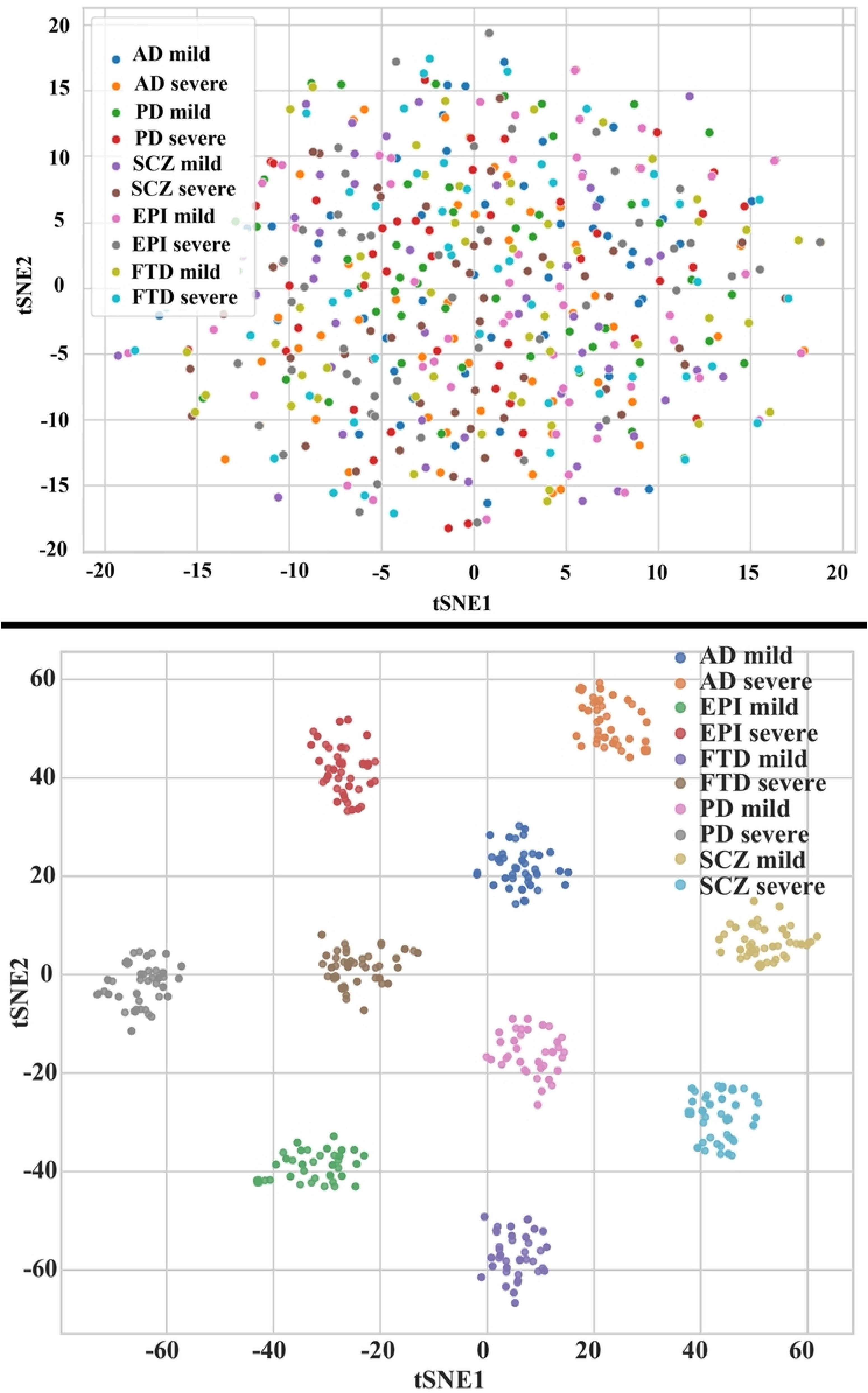
Visualization of penultimate-layer embeddings using t-SNE. A: Embeddings before disease-specific transformation, showing weaker class separation. B: Embeddings after transformation, showing improved clustering and clearer separation between simulated disease conditions.

IG was applied with a zero baseline, and absolute attributions were averaged across the temporal dimension. The resulting per-channel importance profile in Fig 11 shows that the model does not weight all sensors uniformly but concentrates attribution on a subset of electrodes. The learned profile highlights spatial foci consistent with expected pathophysiological priors, including posterior emphasis in AD and motor regions in PD, which grounds model decisions in channel-level neurophysiological structure. The global t-SNE separability indicates inter-class distinctiveness, while the local IG attributions indicate biologically plausible channel contributions.

**Fig 11.**
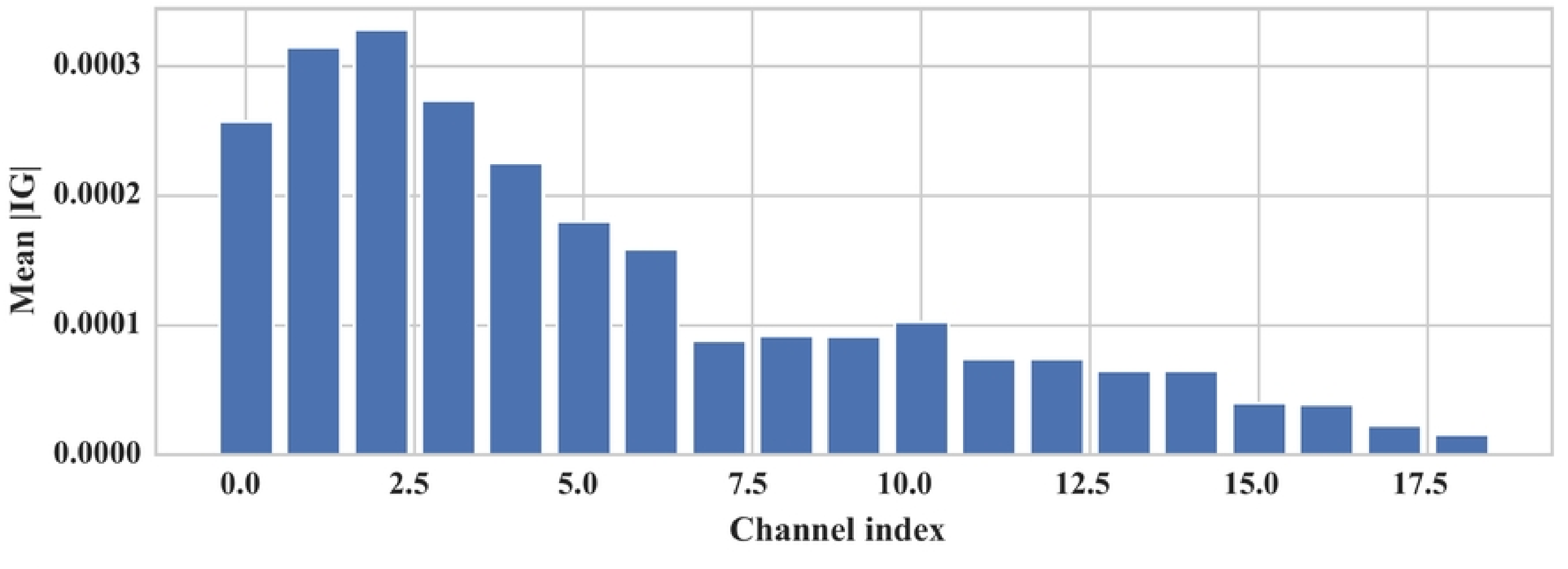
Mean absolute integrated gradients per channel. Attributions are aggregated over time and samples.

## Discussion

NeuroSim emulates clinically relevant neural patterns to support translational neurodiagnostic research. Disease-specific biomarkers, including posterior alpha suppression in AD, exaggerated beta activity in PD, theta and gamma imbalance in SCZ, sharp transients in EPI, and frontal theta escalation in FTD, are integrated systematically and mirror electrophysiological features observed in clinical EEG recordings.

The principal technical distinction of NeuroSim is the integration of multiple clinically motivated controls within a single simulation framework. The framework supports five neurologically distinct conditions within a common signal-generation architecture, whereas many existing approaches are optimized for a single disease, task, or recording paradigm. Disease alterations are explicitly localized to clinically relevant electrode groups rather than applied uniformly across the scalp. The magnitude and manifestation of the injected abnormalities are controlled through severity-dependent parameters, which allows systematic generation of mild, moderate, and severe variants while retaining stochastic subject-level and trial-level variability. The explicit biomarker formulation also provides greater interpretability than purely data-driven generators in which pathological characteristics are learned implicitly.

The primary practical implication is utility in machine learning pipelines where real patient EEG is scarce or privacy sensitive [35]. The augmentation experiment provides direct evidence for this role, since disease-informed synthetic data improved classification of unseen real recordings beyond conventional signal-level augmentation for both AD and PD. Control over disease severity and spatial localization offers further opportunities to investigate differential diagnosis, monitor simulated progression, and evaluate intervention effects, including pharmacological or neuromodulatory treatments [29, 34]. Mahalanobis distance and MANOVA analyses demonstrate that inter-disease differences are statistically more pronounced than intra-disease severity variations, consistent with clinical observations in which diagnostic boundaries outweigh subtle severity gradations [3, 38].

The accessibility of the interface also supports medical education, allowing students and clinicians to explore disorder-specific EEG changes interactively. Beyond diagnostic research, the framework supports BCI development by supplying configurable, noise-controlled datasets for MI, attention decoding, and cognitive-load tasks [31], and it addresses class imbalance for under-represented disorders such as early-stage PD and FTD [38]. Future extensions may incorporate multimodal integration [27] or patient-specific modeling from electronic health records.

### Limitations

The experimentally demonstrated capabilities of NeuroSim are limited to controlled multi-disorder EEG generation, region-specific and severity-specific pattern injection, spectral and spatial biomarker analysis, statistical characterization, synthetic-data classification benchmarking, subject-level augmentation on two real clinical datasets, and technical application of the analysis pipeline to a limited real clinical EEG subset.

Several limitations should be noted. The framework encodes literature-derived pattern priors rather than patient-specific pathophysiology, so simulated signals cannot substitute for clinical recordings in diagnostic validation. The classification benchmark on simulated data is partially circular, as discussed above. Severity levels do not produce strong multivariate separation, which is realistic but limits the use of the framework for severity-grading studies. The TUH EEG analysis uses a small publicly available subset and does not constitute a well-characterized independent diagnostic cohort, because subject-level clinical labels, demographic stratification, and a prespecified clinical endpoint were not included. NeuroSim has not been compared directly with established EEG simulation approaches under an identical downstream task, nor has its effect on a clinical workflow been evaluated with neurologists, EEG technologists, or other intended users. Potential uses in clinical decision support, robustness testing across patient populations, and prospective deployment therefore represent downstream applications that require dedicated validation. Future work should include labeled independent patient cohorts, direct synthetic-versus-real distributional comparisons, established simulator baselines, and prospective evaluation with intended clinical users.

## Conclusion

This study presents a biologically grounded framework for generating synthetic EEG that emulates characteristic patterns of AD, PD, SCZ, EPI, and FTD. Across time-frequency analysis, topographic mapping, and statistical assessment using MANOVA and Mahalanobis distance, the simulated signals align with clinically observed patterns, while an interactive interface supports research prototyping and training. Benchmarking with a CNN-attention model, together with explainability analyses, indicates that the injected patterns are learnable and that model attributions concentrate on pathophysiologically expected electrodes. Subject-level augmentation experiments on two real clinical datasets show that disease-informed synthetic data improve classification of unseen real recordings beyond conventional augmentation strategies. Larger labeled patient cohorts and prospective clinical-user evaluation remain necessary before claims of diagnostic utility or clinical deployment can be made. Future work will extend the engine to multimodal neuroimaging such as combined EEG and functional near-infrared spectroscopy [4, 6, 13, 27, 39], task-specific neural dynamics [17] in further disorders such as autism [40], and patient-specific priors [30].

## Acknowledgments

The real-world clinical EEG data used in this study were obtained with permission from the Neural Engineering Data Consortium at Temple University, specifically the Temple University Hospital EEG Corpus version 2.0.2 [36]. The authors thank the investigators who contributed the data and the funding sources supporting the consortium.

## Data availability

The NeuroSim source code is publicly available at https://github.com/yiamcb/NeuroSim. The clinical EEG datasets analyzed in this study are third-party datasets available from their original providers: the Temple University Hospital EEG Corpus [36], the dataset of Miltiadous et al. [35], and the dataset of Jackson et al. [37].

